# Increased oral Epstein-Barr virus shedding with HIV-1 co-infection is due to a combination of B cell activation and impaired cellular immune control

**DOI:** 10.1101/587063

**Authors:** Catherine M. Byrne, Christine Johnston, Jackson Orem, Fred Okuku, Meei-Li Huang, Stacy Selke, Anna Wald, Lawrence Corey, Joshua T. Schiffer, Corey Casper, Daniel Coombs, Soren Gantt

## Abstract

Epstein-Barr virus (EBV) infection is transmitted by saliva and is a major cause of cancer in people living with HIV/AIDS as well as in the general population. To better understand the determinants of oral EBV shedding we evaluated the frequency and quantity of detectable EBV in the saliva in a prospective cohort study of 85 adults in Uganda, half of whom were co-infected with HIV-1. Participants were not receiving antiviral medications, and those with HIV-1 co-infection had a CD4+ T cell count >200 cells/mm^3^. Daily, self-collected oral swabs were collected over a 4-week period. Compared with HIV-1 uninfected participants, co-infected participants had an increased frequency of oral EBV shedding (IRR=1.27, 95% CI=1.10-1.47). To explain why EBV oral shedding is greater in HIV-1 co-infected participants, we developed a stochastic, mechanistic mathematical model that describes the dynamics of EBV, infected cells, and antiviral cellular immune responses within the tonsillar epithelium, and examined parameter-specific differences between individuals of different HIV-1 infection statuses. We fit the model to our observational data using Approximate Bayesian Computation. After fitting, model simulations showed high fidelity to daily oral shedding time-courses and matched key summary statistics. Examination of the model revealed that higher EBV loads in saliva are driven by B cell activation causing EBV lytic replication in the tonsils, in combination with a less effective EBV-specific cellular immune response. Thus, both these factors contribute to higher and more frequent EBV shedding in HIV-1 co-infected individuals compared to HIV-1 uninfected individuals. These conclusions were further validated by modelling daily oral EBV shedding in a 26-participant North American cohort. Our results provide insights into the determinants of EBV shedding and implicate B cell activation to be a potential therapeutic target to reduce EBV replication in HIV-1 co-infected individuals at high risk for EBV-related malignancies.

**Author summary:** Epstein-Barr virus (EBV) is a ubiquitous infection worldwide. Infection with EBV is associated with the development of several kinds of cancer, including B cell lymphoma and nasopharyngeal carcinoma. Rates of EBV replication and disease are higher in individuals who are also infected with HIV-1. HIV-1 infection is associated with increased B cell activation, which is known to induce EBV reactivation, as well as immunodeficiency resulting from loss of T cells. However, whether these factors contribute to higher rates of EBV replication during co-infection, and by how much, was unknown. We analysed oral EBV shedding data in a cohort of adults from Uganda that were chronically infected with EBV. We found that participants that were HIV-1 infected were much more likely to have detectable quantities of EBV in their saliva. Also, when detected, the quantity of EBV present in the saliva was usually higher in HIV-1 infected participants. To better understand these findings, we developed a mathematical model to describe the dynamics of EBV, EBV-infected cells, and the cellular immune response within the tonsils. By rigorously matching our model to our participant data, we determined that high EBV loads in saliva are caused by high rates of infected B cell activation, as well as worse cellular immune control of EBV infection. These results provide an explanation of the impact of HIV-1 on EBV infection. Further, they suggest that strategies that suppress B cell activation may prevent EBV-related malignancy in people who are also infected with HIV-1.

## Introduction

Epstein-Barr virus (EBV) infection is associated with the development of approximately 200,000 malignancies per year, including B cell lymphomas and nasopharyngeal carcinoma [1]. The risk of EBV-associated malignancies is significantly higher in people co-infected with HIV-1. For example, risks of non-Hodgkin lymphoma in the U.S., an AIDS-defining cancer, are 10-fold higher in HIV-1 infected individuals than the general population [2]. Individuals with EBV/HIV-1 co-infection tend to have higher EBV viral loads in saliva and blood [3–5]. Uncovering the mechanisms by which HIV-1 impairs control of EBV infection may provide clues relevant to prevention of EBV-related disease as well as insights into basic EBV pathobiology.

EBV is primarily transmitted via saliva, and infection is nearly universal, occurring during early childhood in developing countries and by young adulthood in highly developed countries [6–8]. During primary infection, it is thought that EBV first infects oral epithelial cells overlying the lymphoid tissue known as Waldeyer’s ring [9]. This area consists of the tubal, adenoid, palatine and lingual tonsils [9]. Infected epithelial cells produce large numbers of infectious virions [10], facilitating latent infection of naïve B cells in the underlying lymphoid tissue. EBV drives these naïve B cells to mature into resting memory B cells and circulate throughout the body, through the expression of only a small number of latent gene products [11, 12]. Viral shedding is highest during primary EBV infection but remains frequent throughout chronic infection [5]. During chronic infection, B cells latently infected with EBV can return to Waldeyer’s ring, encounter cognate antigen, and become activated to mature into plasma cells, triggering lytic reactivation and production of infectious virions [13–15]. This process initiates a new round of epithelial infection in the tonsils and viral shedding in the saliva. Increased EBV shedding with HIV-1 co-infection may be due to more frequent reactivation of EBV-infected B cells, and thus increased viral seeding of oral tissue, and/or impaired T cell-mediated immune control of EBV replication, prolonging or inhibiting the clearance of infected epithelial cells [16–18].

The dynamics of chronic herpesvirus infections in humans have been elucidated by longitudinally sampling mucosal surfaces and blood, revealing the patterns of latency, reactivation, and dissemination, as well as giving insight into viral pathogenesis and host-pathogen interactions [19, 20]. While several studies have examined EBV mucosal shedding patterns in HIV-1-infected persons [5, 21–25], the majority have been in the setting of advanced HIV-1 infection or in persons receiving highly active antiretroviral therapy (HAART) [23–25]. The mechanisms by which co-infection with HIV-1 significantly increases viral loads in saliva have not been explained in part due to the requirement for frequent quantitative measures of shedding amenable to mathematical modelling [23–25]. Previous mathematical models have examined the within-host dynamics of EBV infection, developing quantitative descriptions of how infected cells, B cells, antibodies, virus, and T cells interact, and determining biologically relevant parameter values for the rates at which infection dynamics occur in the tonsillar epithelium [10, 26–30]. However, none have examined the differences between EBV shedding in HIV-1 co-infected and HIV-1 uninfected individuals. In this study, we analysed rich oral EBV shedding data from a cohort of Ugandan adults with or without HIV-1 co-infection and constructed a mechanistic mathematical model to identify the determinants of EBV replication.

## Results

### HIV-1 infection is associated with increased frequency and quantity of oral EBV shedding

Among this cohort of 85 participants, a total of 2264 daily oral swabs were collected, with a median of 29 swabs per participant (range 1-32). 43 (51%) participants were HIV-1 seropositive. Additional details of the cohort have been previously published [31], and other data from the cohort, including EBV loads in genital swabs and plasma samples, can be found in the Supporting Information. We began our analysis by first examining whether HIV-1 infection status affects the frequency of EBV detection in the saliva. We found that HIV-1 infection was significantly associated with EBV detection in oral swabs, increasing the frequency of observation 1.27-fold (CI = 1.10-1.47, p-value = 0.001, Fig 1A) and increasing the median quantity of EBV detected in oral swabs by 1 .61 log_10_ genome copies (CI = 1.29-1.93, p-value <0.001, Fig 1B). We also saw large variability in participant viral loads over time (Fig 1C). In HIV-1 uninfected participants, the median percentage of swabs positive for EBV was 86% (range 0-100%, interquartile range (IQR) 33%) while in HIV-1 co-infected participants the median percentage of swabs positive for EBV was 100% (range 27-100%, IQR 0%). Of swabs that tested positive for EBV, viral loads varied over time by a median of 3.49 orders of magnitude (range 0.34-5.27, IQR 1.32) within individual HIV-1 uninfected participants and a median of 2.30 orders of magnitude (range 0.95-5.93, IQR 1.15) within individual HIV-1 co-infected participants.

**Fig 1.**
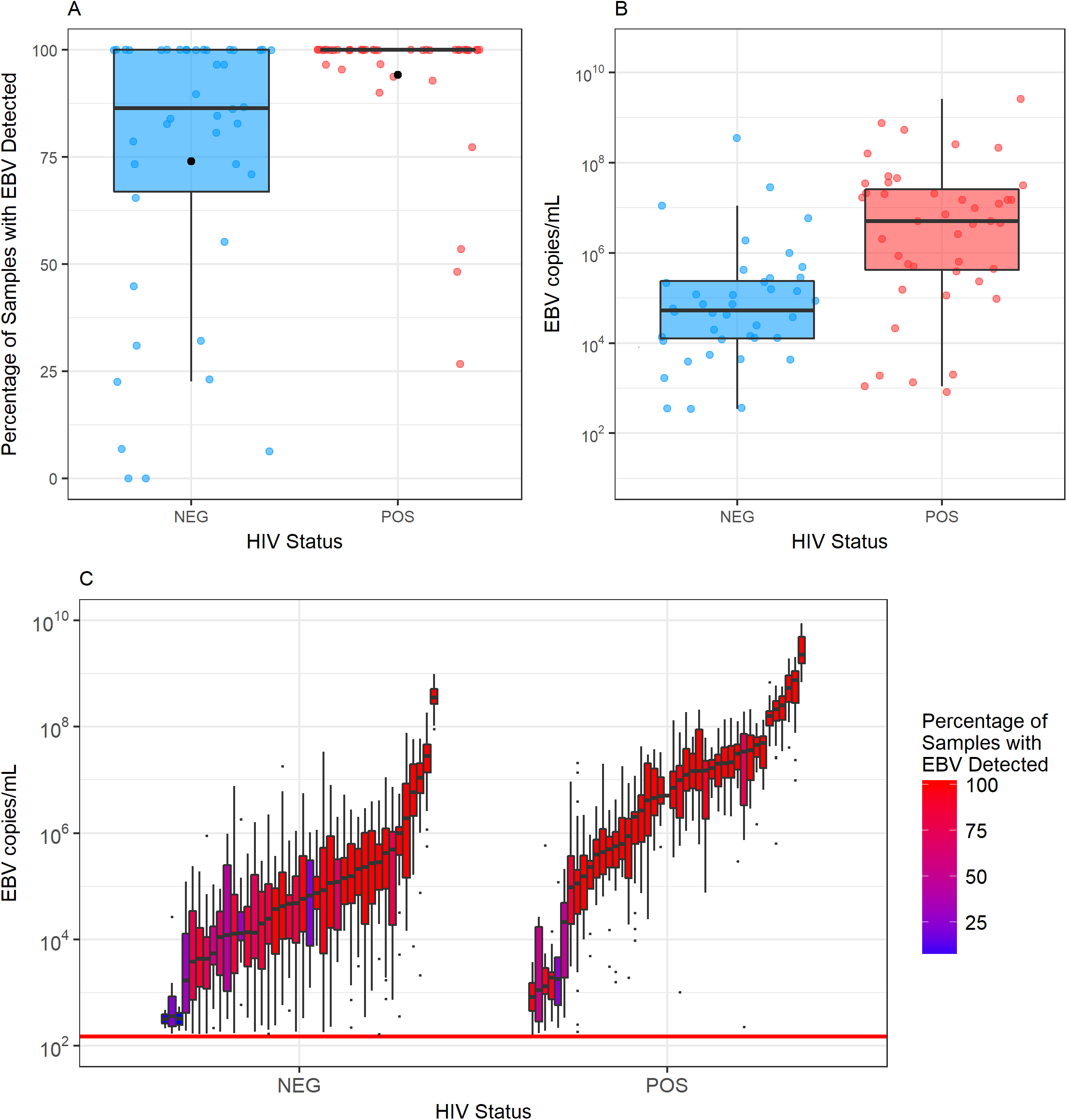
Impact of HIV-1 infection on EBV detection A. Percentages of saliva samples that tested positive for EBV for each participant stratified by HIV-1 status. Black dots indicate the percentage of samples that tested positive for EBV when pooling participant samples. B. Median EBV viral loads/ml in oral swabs testing positive for EBV for each participant stratified by HIV-1 status. C. Distributions of participants’ oral swab viral loads. Each box and whisker represents the viral loads of EBV-positive oral swabs for an individual participant. The percent of oral swabs that tested positive for EBV for each participant is indicated by the colour of the box. The red horizontal line represents the threshold of EBV detection (150 EBV copies/ml).

### The effect of CD4+ T count and HIV-1 plasma RNA levels on oral EBV shedding in HIV-1 co-infected individuals

We next analysed how CD4+ T cell counts and HIV-1 plasma RNA levels impacted the frequency of EBV detection in oral swabs of HIV-1 infected participants (Table 1). CD4+ T cell count significantly affected the frequency of EBV detection in oral swabs. Each additional 100-cell increase in CD4+ T cells/mm^3^ was associated with a 5% reduction in the odds of EBV detection. This is consistent with cell-mediated immunity conferring partial but incomplete control of EBV replication. We found no significant association between HIV-1 viral load in plasma and the frequency of EBV detection in saliva.

**Table 1.** Effects of plasma HIV-1 load and CD4+ T cell count on the frequency of EBV detection. The effects of each 100-cell increase in CD4+ T cells/mm^3^ and each log_10_ increase in HIV-1 RNA on the incidence rate ratio (IRR) of EBV detection, their confidence intervals (CI) and their p-values are shown.

| Trait | IRR | 95% CI | p-value |
| --- | --- | --- | --- |
| HIV-1 RNA | 1.04 | 0.98-1.12 | 0.201 |
| CD4+ T cell | 0.95 | 0.92-0.99 | 0.008 |

### Mathematical Model of EBV Shedding in the Tonsils

To obtain mechanistic insights into chronic oral EBV shedding and to better understand the drivers of lytic replication and transmission, we constructed a mathematical model that captures the relevant anatomic, virologic and immunologic features of oral EBV infection. In chronically infected individuals, EBV is shed in all areas of Waldeyer’s ring including the palatine, lingual, tubal tonsils and adenoids [9]. Most of the tonsillar area is composed of stratified squamous epithelium or ciliated pseudostratified columnar epithelium, arranged into a series of crypts or folds, allowing for a large surface area [32]. The epithelium is often only one cell thick where EBV can transcytose to reach the underlying lymphoid tissue where B cells and germinal centres are found [33]. The palatine tonsils have an estimated surface area of 295 cm^2^, arranged into approximately 20 crypts [32], while the lingual tonsil area is composed of 35-100 crypts, and the adenoids are composed of a series of folds in lymphoid tissue [34]. By estimating that each palatine tonsil is approximately 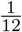 of the entire surface area of Waldeyer’s ring, a series of 240 crypts, each serving as sites where EBV infection may occur, can represent the entire region. We assumed that the dynamics of each crypt are independent from each other and explicitly modelled the dynamics of infected epithelial cells, the immune response, and viral load within each crypt.

The dynamics within each crypt are shown in Fig 2. All parameters were assumed to remain constant over the course of study, with some varying between participants to account for differences in EBV shedding and HIV-1 infection status. We assumed that latently infected B cells reactivate and infect the epithelium at a constant rate *b*. Infected epithelial cells, *I*, infect other epithelial cells through cell-to-cell contact at a constant per-capita rate *β*. As the total number of cells within a single crypt is large in comparison to the expected number that can become infected with EBV, we assumed that target cell number is not limiting. Epithelial infection causes the recruitment and proliferation of EBV-specific cytotoxic T cells, *T*, at a per-capita rate *θI*. Cytotoxic T cells kill infected epithelial cells following the law of mass action at a rate *fIT*. We assumed that cytotoxic T cells die or leave the tonsils at a per-capita rate *δ*. Independent of infection, we assumed a constant number of EBV-specific cytotoxic T cells, *α*, are tissue-resident. Like *T*, these cells can kill infected epithelial cells and stimulate the proliferation of new EBV-specific cytotoxic T cells; however, while population *T* leaves the system over time, these tissue-resident T cells remain within the tissue and do not recirculate. This means there are always immune cells present to respond to new infection and tissue is never entirely unprotected, allowing for faster control of infection. EBV virions, *V*, are produced by infected epithelial cells, enter saliva for at a per-capita rate *p* and are cleared at a per-capita rate *c*. In this model, we assumed the main contributors to virus in the saliva are infected epithelial cells and thus we did not directly model virions produced by infected B cells [10]. With the propagation of EBV infection shown to be 800-fold more efficient through cell-to-cell contact rather than through free virus, we also chose to assume all new epithelial cell infection is caused by cell-to-cell contact [10, 35].

**Fig 2.**
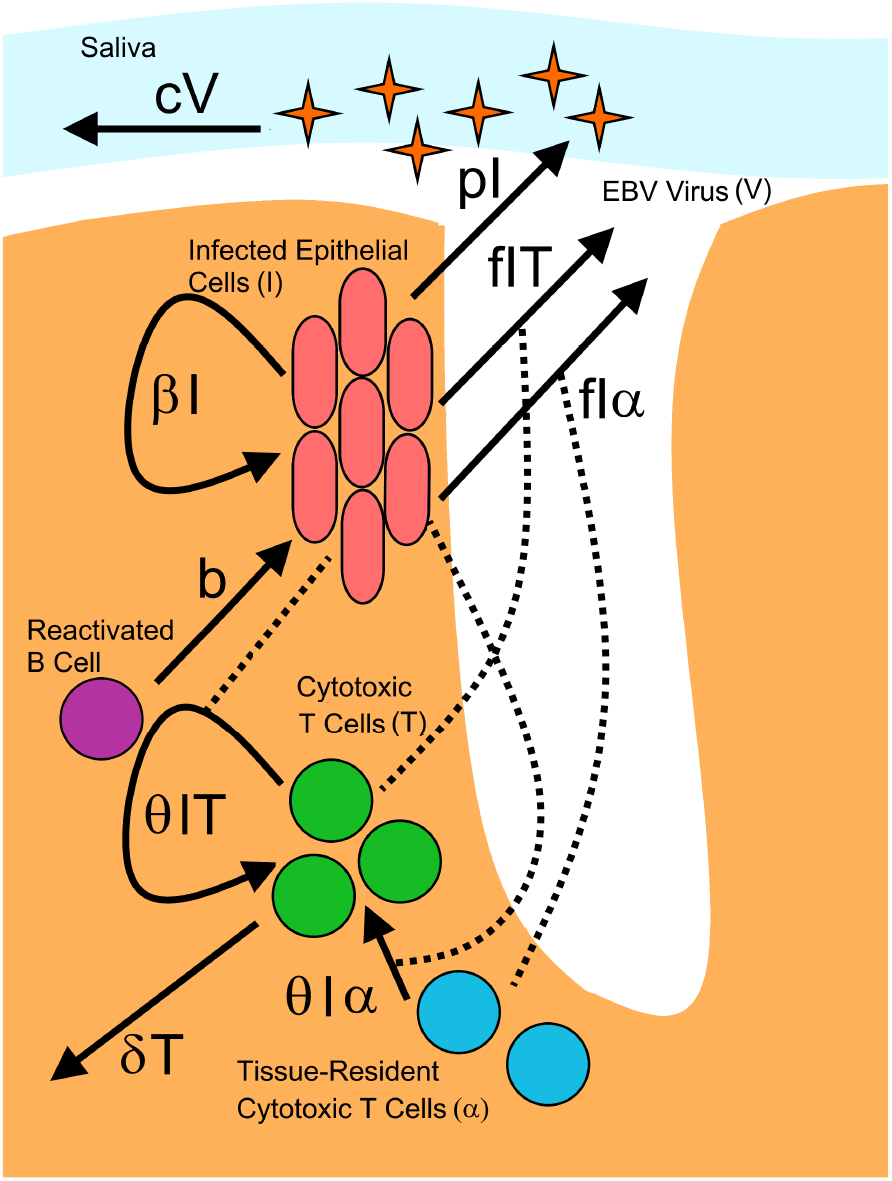
Description of single crypt dynamics. Waldeyer’s ring is represented as a series of 240 individual crypts in which infection dynamics occur. Within each crypt, the population dynamics of infected epithelial cells (I), cytotoxic T cells (T) and EBV (V) are described. The salivary viral load is represented by the total viral load aggregated across all crypts.

The concentration of EBV detected in the saliva of participants was highly variable, and frequently undetectable. Therefore, we chose to implement our model in a stochastic framework in order to capture these traits (Methods). Our model assumptions were used to build a chemical master equation system [36] that describes all system reactions within a single crypt, as follows:

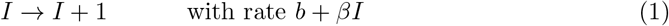

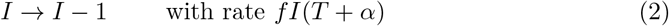

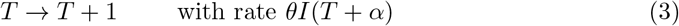

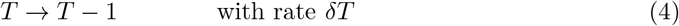

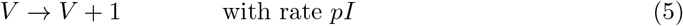

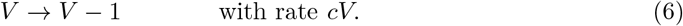

### Basic model analysis

To understand how the different parameters of our mathematical model affect simulated EBV shedding in saliva, we first conducted a literature review to find previously estimated or measured values of our parameters (Table 2). We then performed a simple univariate analysis to evaluate how changes in these parameters influence the characteristics of model simulations. The starting set of parameters was chosen based on current estimates in the literature; we then performed multiple model simulations, varying each parameter in turn over 4 orders of magnitude to observe how sensitive the model is to these changes (Fig 3). Since we observed large variability in the viral loads of different cohort participants (Fig 1C), our aim was to select a fixed set of parameters that would allow for the generation of a wide range of viral loads. Increases in parameter *b* or *δ* increased the median viral load of our model simulations; however, the maximum viral load remained stable. In contrast, *β* appeared to control the maximum viral load reached in simulations, with higher *β* values causing higher viral loads and larger variance in the viral load. Parameters *f, α*, and *θ*, all governing the strength of the cellular immune response, appeared to have similar effects on the simulation, all causing comparable decreases in viral loads as the magnitude of the parameter increased. Intuitively, increases in parameter *p*, governing the viral burst size, increased the viral loads in simulations. Low viral loads and periods of viral extinction (as seen in many HIV-1 uninfected participants) were uncommon with higher values of *p*. Thus, these higher values of *p* are improbable. Lastly, increases in parameter *c* reduced the viral loads and increased the variance. From this analysis, we chose our fixed parameter values to be *β* = 50, *f* = 0.1, *α* = 200, *δ* = 0.1, *p* = 10^4^, and *c* = 6, which generally agreed with published estimates. We allowed *b* and *θ* to vary during fitting as these two parameters are the most likely to be affected by HIV-1 infection status.

**Fig 3.**
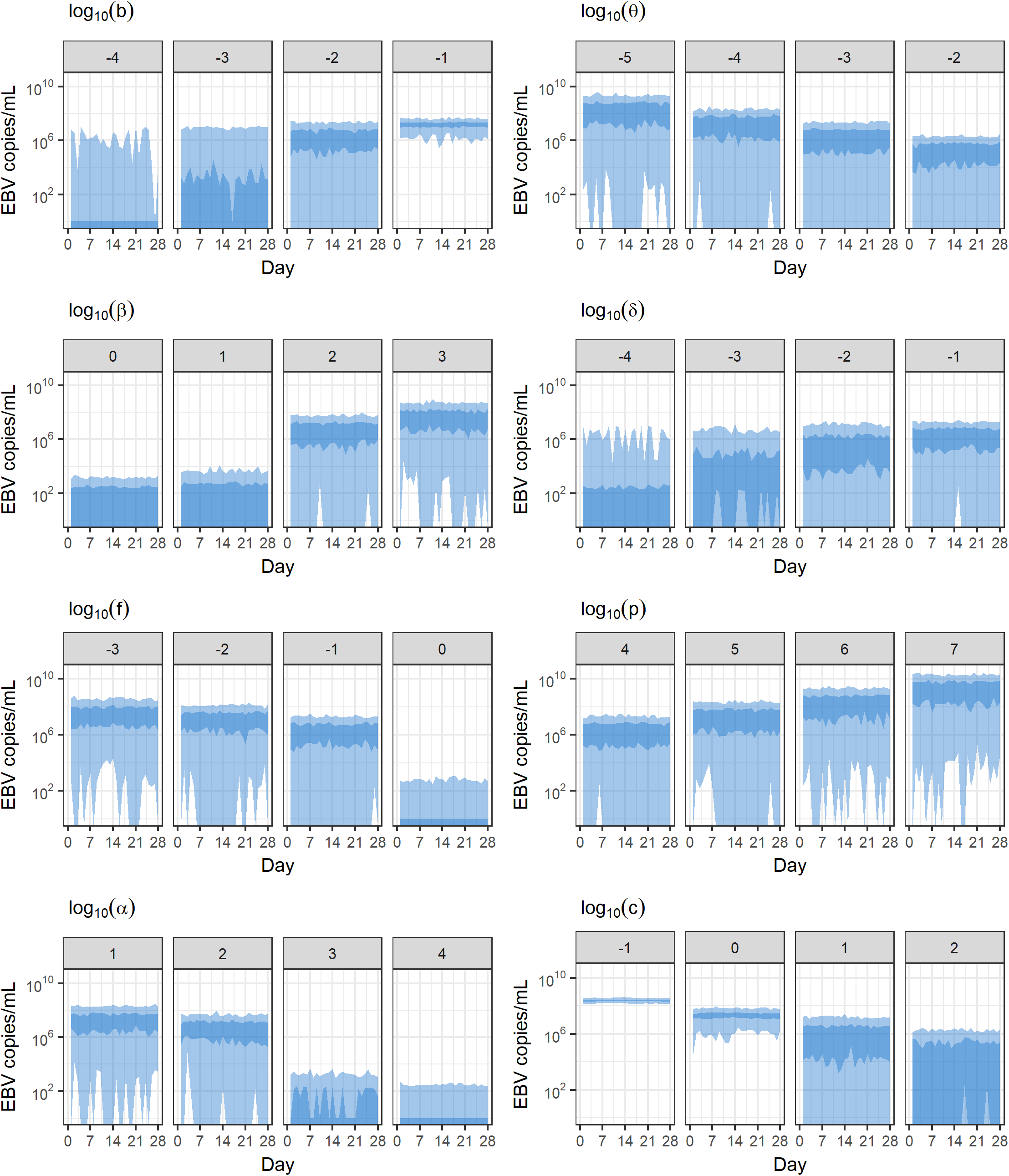
Evaluation of individual model parameters. Each parameter of the model was examined to determine how it influences the simulation of oral viral shedding over time. Parameters were varied one at a time over 4 orders of magnitude while keeping all others constant, and 100 simulations were run for each parameter set. Viral quantity over time is shown; 25-75% quartiles are shown in dark blue while 0-100% quartiles are show in light blue. When not varying, *b* = 0.01, *β* = 50, *f* = 0.1, *α* = 200, *θ* = 0.001, *δ* = 0.1, *p* = 10^4^, and *c* = 6.

**Table 2.** Parameter values used in the model. Parameters that remained fixed throughout cohort data fitting were chosen based on published values and univariate analysis. * The discrepancy between the published and model values for parameter *β* is due to previous models separately accounting for infection due to cell-free virus. ** The discrepancy between the published and model values for parameter *δ* is due to previous models not accounting for tissue-resident cytotoxic T cells as a separate population.

| Parameter | Units | Description | Published Values | Model Values |
| --- | --- | --- | --- | --- |
| $b$ | cell day<sup>-1</sup> | rate of B cell reactivation causing new lytic epithelial infection | - | fitted, see text |
| $\beta$ | day<sup>-1</sup> | per-capita rate of cell-to-cell infection | 1.6 [37] | 50* |
| $f$ | day<sup>-1</sup> cell<sup>-1</sup> | per-capita death rate of infected epithelial cells due to the effect of cytotoxic T cells | $1.8 \times 10^{-9} - 0.2$ [27, 38, 39] | 0.1 |
| $\alpha$ | cell | number of tissue-resident cytotoxic T cells | - | 200 |
| $\theta$ | day<sup>-1</sup> cell<sup>-1</sup> | per-capita proliferation rate of cytotoxic T cells dependent on the number of infected epithelial cells | - | fitted, see text |
| $\delta$ | day<sup>-1</sup> | per-capita death rate of cytotoxic T cells | $9.5 \times 10^{-3}$ [27, 40] | 0.1** |
| $p$ | virion day<sup>-1</sup> ml<sup>-1</sup> cell<sup>-1</sup> | production rate of EBV virions by infected epithelial cells | $10^2 - 10^5$ [10, 27] | $10^4$ |
| $c$ | day<sup>-1</sup> | per-capita clearance rate of EBV virions | 2 [27, 41] | 6 |

### Mathematical model fits clinical data well and simulates oral shedding data with high fidelity

We fit our mathematical model to each participant’s data using Approximate Bayesian Computation (ABC). As our model is stochastic, this fitting process involved finding parameter values that produced simulations with similar summary statistics to the data rather than simulations that directly matched the curve of the data. The traits of each participant were captured using five summary statistics: the percentage of EBV-positive swabs, the median, maximum, and variance of detectable viral loads, and the number of peaks in viral load, a peak being defined as when the directly preceding and following time points have lower viral loads. The goodness of fit was assessed by the statistic *ρ_i,j_*, which is defined as

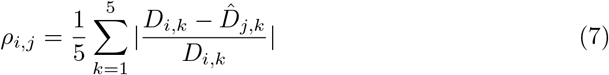

for participant *i* and parameter set *j*. Here *D_i,k_* is the *k^th^* summary statistic for the data of participant *i* and 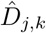 is the *k^th^* summary statistic for simulations using parameter set *j*. Lower *ρ* values indicate a better fit between the model simulation and the data. Full details are given in the Methods. Examples of 4 participants’ shedding data and model simulations with *ρ* values varying between 0.1 and 0.7 are shown in Fig 4. At these low *ρ* values, all simulations capture the summary statistics of the participants quite well, but a general improvement in fit quality is apparent as *ρ* approaches 0.1.

**Fig 4.**
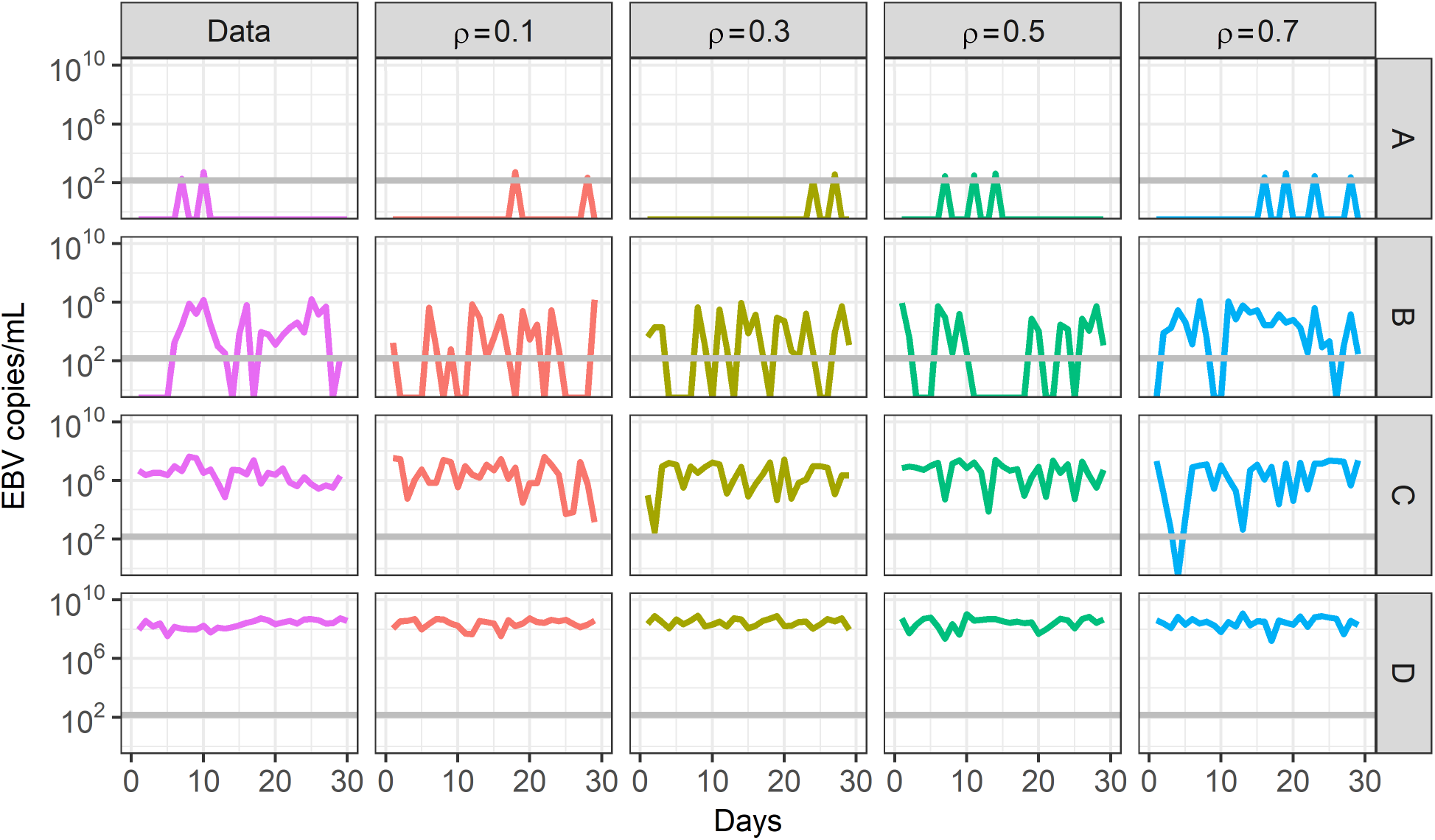
Comparison of EBV shedding patterns in Ugandan cohort participants and model simulations. The EBV shedding patterns of four representative participants are shown. Participants A and B are uninfected with HIV-1 while participants C and D are co-infected with HIV-1. Model simulations that fit these data with varying success (*ρ* = 0.1, 0.3, 0.5, 0.7) are shown. Lower *ρ* values indicate a better fit to the summary statistics of the data. The horizontal grey line indicates the threshold of detection (150 EBV copies/ml) when measuring EBV loads in participant saliva samples.

Among all 85 participants’ data, we were able to fit parameters to 82. Of the 3 participants whose data could not be fit, 2 participants had no EBV detected in any of their saliva swabs, and 1 participant had only 1 swab taken. Fig 5 shows the distribution of *ρ* values (Equation 7) calculated for 1000 optimal parameter sets for each participant that was fit by the model. The *ρ* values accepted during fitting ranged between 0.002 and 0.558, indicating that accepted model fits matched the data at least as well as the examples shown in Fig 4. In general, our model fits were slightly better for participants with a high median viral load. Using generalized estimating equations (GEE), and assuming a Gaussian distribution for *ρ* values, we found that each log_10_ decrease in a participant’s median viral load increased *ρ* by 0.051 (95% CI =0.039-0.063, p-value< 0.001).

**Fig 5.**
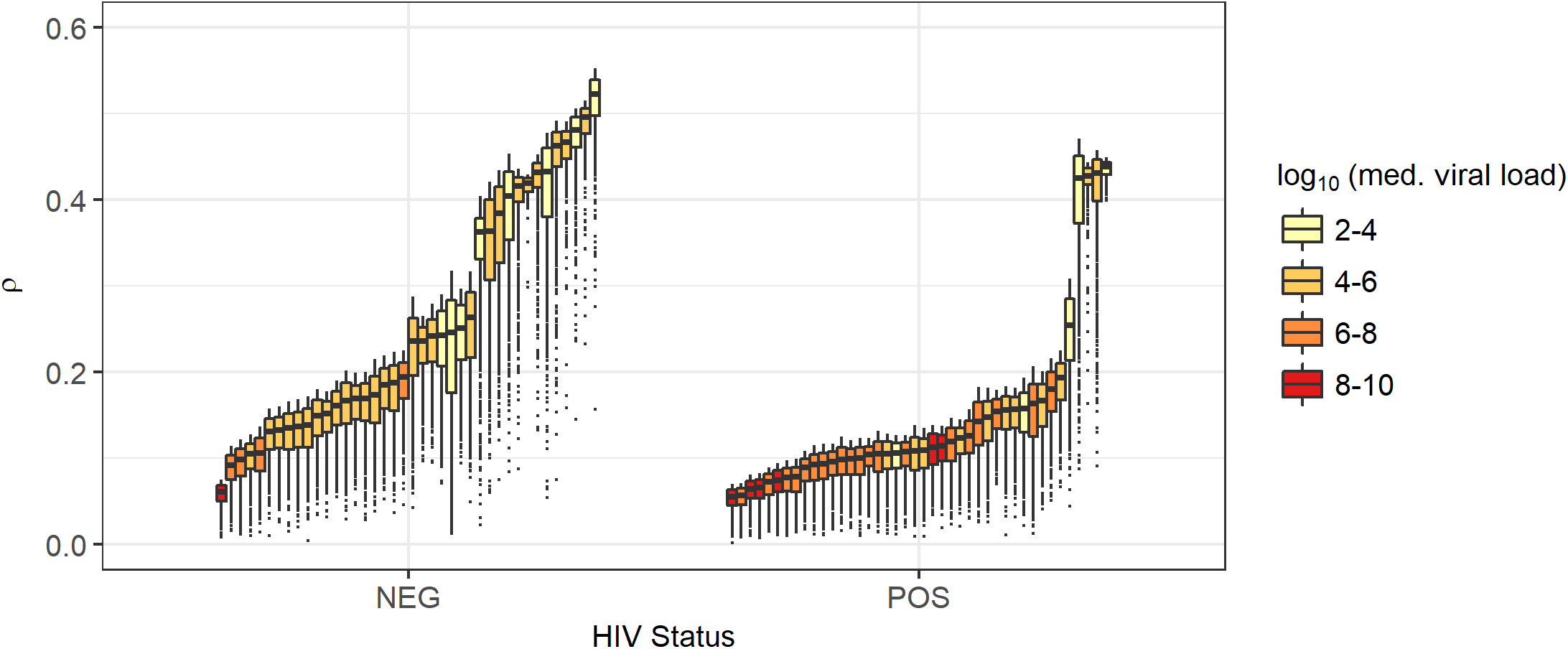
Goodness of fit of mathematical model to participant data. Datasets from 82 study participants were fit by our mathematical model. Each participant’s dataset is represented by 1000 parameter sets that best fit model simulations to the data. Goodness of fit for each parameter set is measured by a *ρ* value as defined in Equation 7. Boxes indicate the interquartile range and whiskers indicate the 95% range. Lower values indicate a better fit between the data and the model. The colour of the box indicates the median EBV viral load detected in the saliva of that participant.

### Sensitivity analysis of the fitted model

After fitting our model to data (Methods), we performed a sensitivity analysis to confirm that we chose suitable values for the parameters that remained fixed throughout the fitting process. Parameters *b* and *θ* were fixed at their fit values, while those that had remained fixed during fitting were varied over two orders of magnitude. A simulation was run for each new set of parameters and the goodness of fit was compared to the original. Results are shown in Fig 6. In all cases and for all participants, the median *ρ* value generated from parameter sets was higher and, therefore, a worse fit than our fixed parameter choices, providing confidence that the values chosen for our fixed parameters provided a consistently good fit.

**Fig 6.**
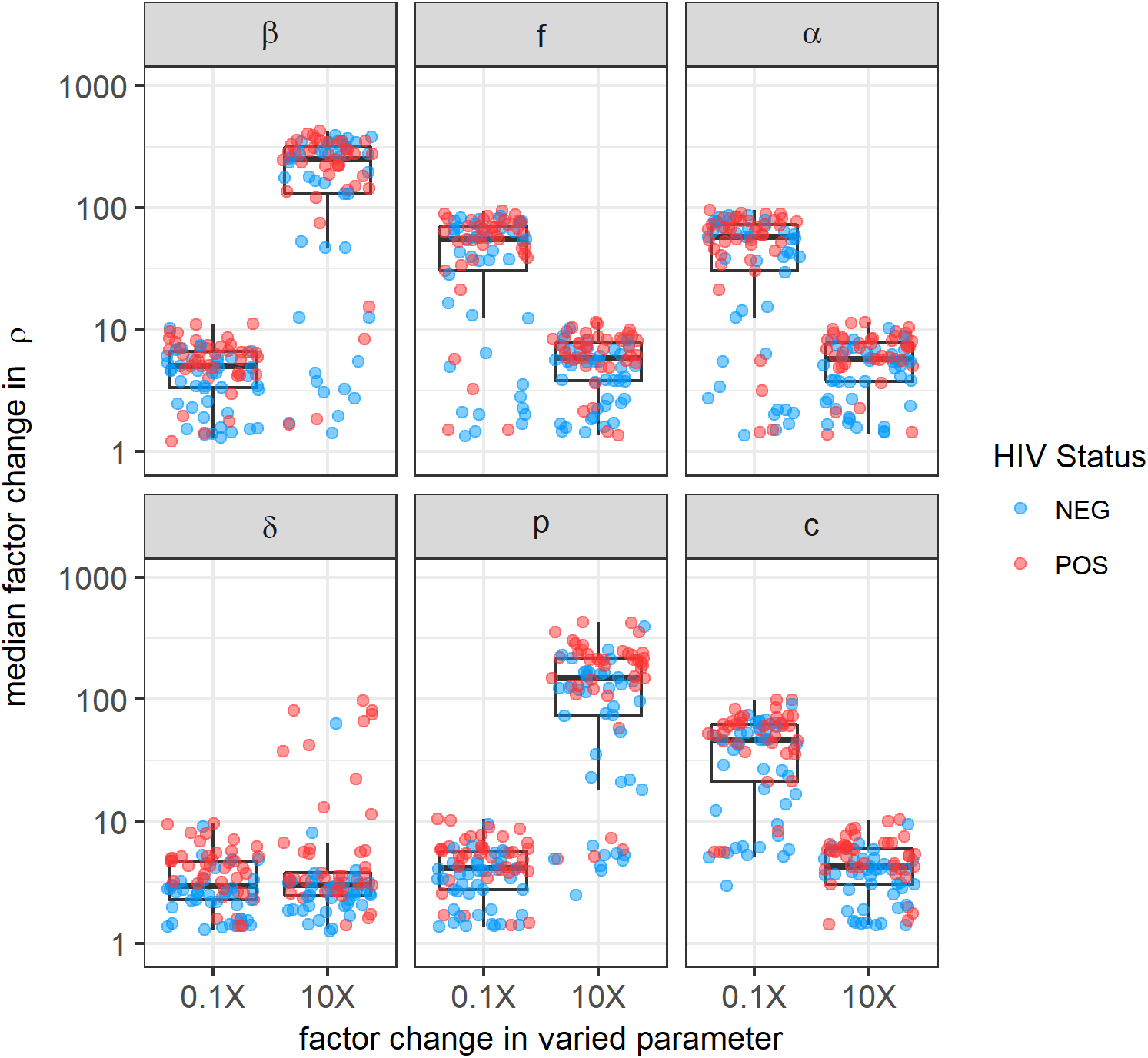
Sensitivity analysis of fit parameters. For the 1000 parameter sets selected by the ABC fitting algorithm for each participant, we varied parameters *β, f, α, δ, p*, and *c* (the parameters that remained fixed during fitting) over two orders of magnitude to observe whether changes in these values could have improved the model fit. Each parameter was set to equal 0.1 and 10 times the value it was set to during fitting and one simulation of each new parameter set was performed. The median factor-change in the resulting *ρ* value for each participant is shown on the y-axis. In all cases, the new parameter values led to worse fits (factor change in *ρ* exceeded one).

### High EBV viral loads are caused by large numbers of actively infected crypts

When examining the model-predicted viral dynamics in simulated individuals, we noticed that high viral loads in the saliva were caused by multiple crypts actively producing virus at the same time (Fig 7). Using importance sampling (Methods) on the results of our ABC fitting algorithm, we achieved a representation of how the number of actively infected crypts and the amount of virus produced by these crypts differ among individuals stratified by HIV-1 infection status.

**Fig 7.**
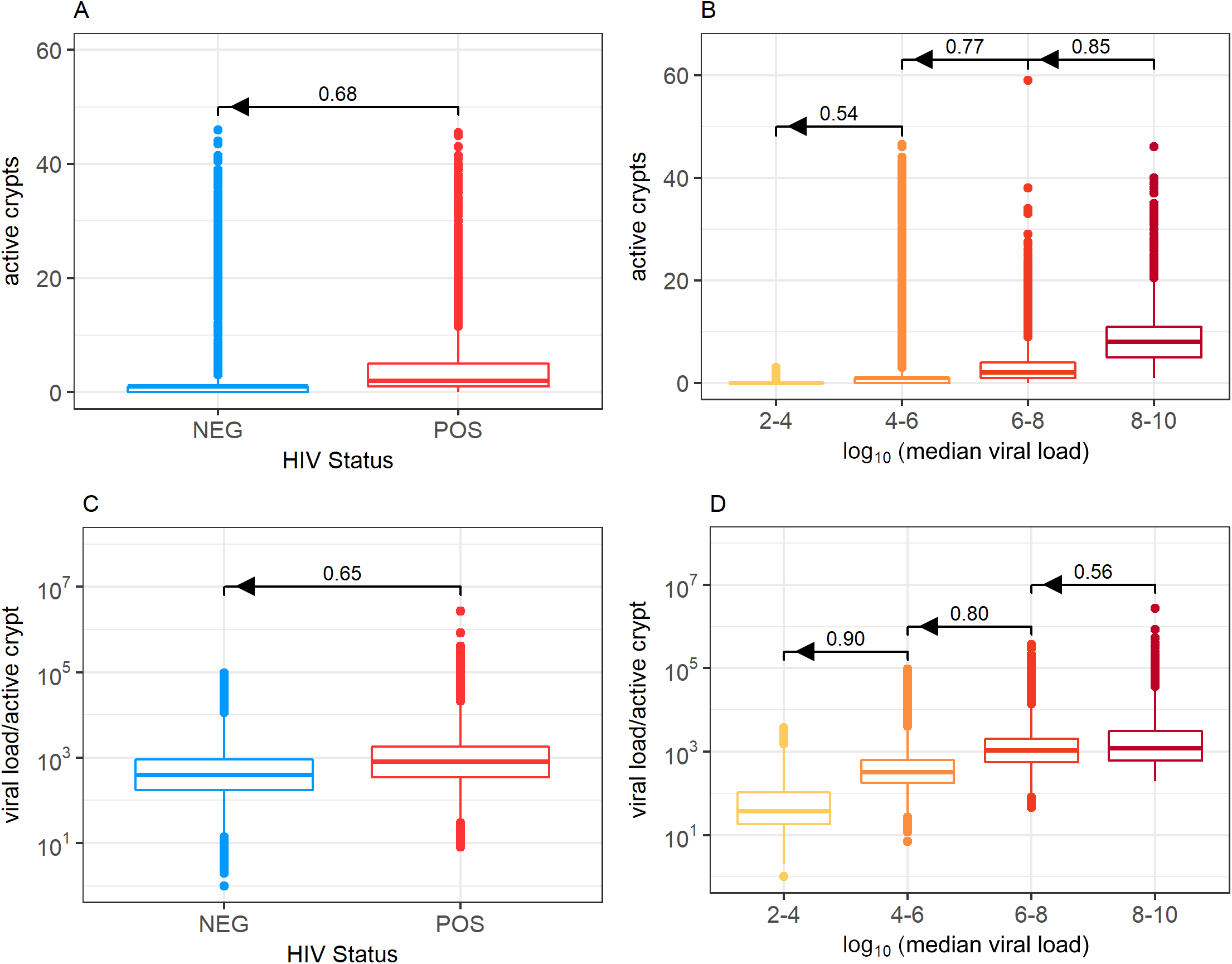
Predicted numbers of active crypts and viral load per active crypt. Distributions of the median number of crypts actively producing EBV (A and B) and the median EBV viral load produced by an active crypt at any given time (C and D) are shown stratified by participant HIV-1 status and EBV median viral load in the saliva. Increases in median salivary EBV viral load are caused by a higher number of crypts having active (B), and more extensive (D) infection. This trend translates to HIV-1 co-infected participants having more infected crypts infected, and each infected crypt producing more virus. Bars above boxplots indicate the probability that a randomly selected individual of one group has a higher parameter value (be it number of active crypts or viral load/active crypt) than a randomly selected individual in a second group. Arrows show the direction of comparison. We see that HIV-1 uninfected individuals usually have more actively infected crypts and more virus per active crypt than HIV-1 co-infected individuals. Similarly, we see that individuals with higher median viral loads in their saliva usually have more actively infected crypts and more virus per active crypt than individuals with lower median viral loads.

The distributions for the median number of crypts HIV-1 co-infected and HIV-1 uninfected individuals have over time are shown in Fig 7A. Over time, an HIV-1 co-infected person will have a higher median number of actively infected crypts than an HIV-1 uninfected person with probability 0.68 (Fig 7A). While these distributions vary greatly (IQR of 1 and 4 for HIV-1 uninfected and HIV-1 co-infected individuals respectively), over time HIV-1 uninfected persons are expected to have a median of 1 crypt within their tonsils actively producing virus, while HIV-1 co-infected individuals are expected to have 2. These results match well with previous estimates that indicate individuals have ≤ 3 independent plaques of oral epithelial infection at any given time [10]. Infected crypts within an HIV-1 infected individual are also shown to produce more virus. Over time, an HIV-1 co-infected individual is expected to have a higher median viral load within their actively infected crypts than an HIV-1 uninfected individual with probability 0.65 (Fig 7C). Our simulations predict a median of 798 EBV DNA copies per active crypt in HIV-1 co-infected individuals and 389 EBV DNA copies per active crypt in HIV-1 uninfected individuals. These distributions again vary greatly, leading to overlap (IQR of 753 and 1472 for HIV-1 uninfected and HIV-1 co-infected individuals respectively). The overlap between the distributions of traits for HIV-1 co-infected and HIV-1 uninfected individuals implies that HIV-1 status alone does not determine a distinct phenotype. Thus, we chose to also examine the behaviour of traits when stratifying our simulations according to participants’ median viral loads in the saliva. Higher median EBV viral loads detected in saliva correlate well with higher numbers of actively infected crypts (Fig 7B) and higher viral loads per actively infected crypt (Fig 7D). Together, these results indicate that high EBV loads in the saliva are caused by more frequent and extensive infection in tonsillar crypts.

### Greater oral EBV shedding in HIV-1 co-infection is due to increased B cell reactivation and weaker cellular immune response

We next looked at parameter-specific differences between individuals of different infection status groups and determined distributions for parameters *b* (rate of B cell reactivation causing new lytic epithelial infection) and *θ* (rate of EBV-specific cytotoxic T cell proliferation and recruitment). Parameter distributions stratified by HIV-1 infection status and median EBV viral load in saliva are shown in Fig 8. While these distributions display substantial variability and overlap, changes in both *b* and *θ* account for the the generally higher EBV viral loads observed in the saliva of HIV-1 co-infected individuals. HIV-1 co-infected individuals appear to have higher rates of B cell reactivation. The median value of *b* was found to be 2.9 times higher in HIV-1 co-infected individuals, equalling 0.007/day in HIV-1 uninfected individuals (IQR for *b* of 0.01) and 0.02/day in HIV-1 co-infected individuals (IQR for *b* of 0.04). Together, these distributions indicate that an HIV-1 co-infected individual will have a higher *b* value than an HIV-1 uninfected individual with probability 0.76. HIV-1 co-infected individuals also appear to have lower rates of EBV-specific cytotoxic T cell immune response. The median value of *θ* was 19.7 times lower in HIV-1 co-infected individuals, equalling 3.1 × 10^−4^ in HIV-1 co-infected individuals (IQR for *θ* of 0.04) and 6.1 × 10^−3^ in HIV-1 uninfected individuals (IQR for *θ* of 0.001). Together these distributions indicate that an HIV-1 co-infected individual will have a lower *θ* value than an HIV-1 uninfected individual with probability 0.74. We also note that, in general, increases in median EBV viral load in saliva correlate with increases in the rate at which B cell reactivation occurs and causes infection (*b*). Similarly, there is an inverse correlation between the strength of the EBV-specific cytotoxic T cell immune response (*θ*) and the median viral load (Fig 8B and 8D).

**Fig 8.**
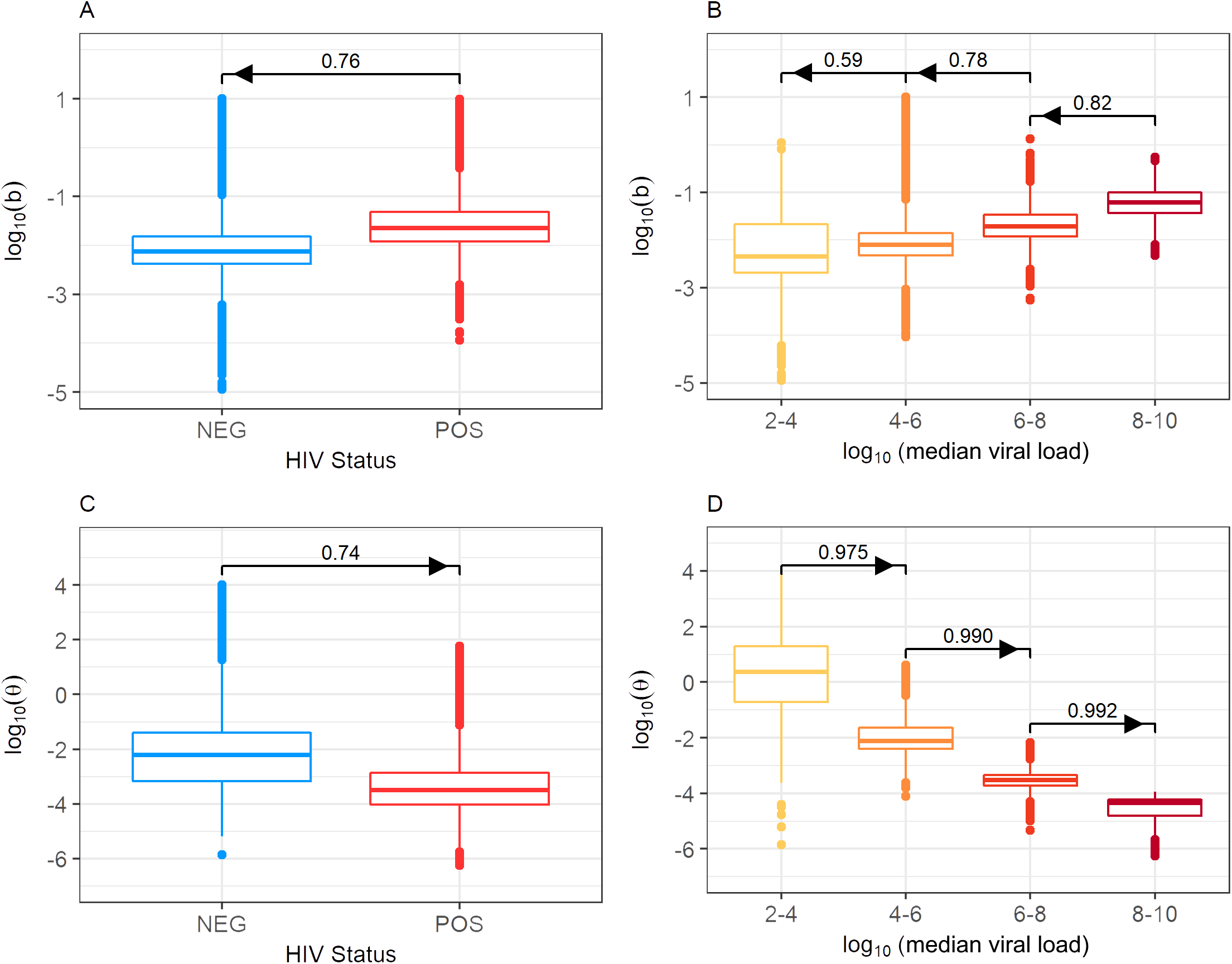
Distribution of parameters *b* and *θ*, stratified by HIV-1 status and median EBV viral load in saliva. Fitting our mathematical model to participant data revealed that parameter *b* is usually greater in HIV-1 co-infected participants (A) and increases with median EBV viral load (B), and parameter *θ* is usually lower in HIV-1 co-infected participants (C) and decreases with median saliva EBV viral load (D). Bars above boxplots indicate the probability that a randomly selected individual of one group has a higher parameter value (be it *b* or *θ*) than a randomly selected individual in a second group. Arrows show the direction of comparison.

We also sought to quantify the correlation between *b* and *θ* (Fig 9). When looking at within-group correlation of participants with similar median viral loads, *b* and *θ* have moderate positive correlation (R = 0.22, 0.42, 0.36 and 0.71 for increasing viral load groups), indicating that *b* and *θ* can counter-balance to cause similar viral loads. However, when observing all data, *b* and *θ* are moderately negatively correlated (R = 0.45), indicating the highest viral loads are found in individuals whose parameters feature high *b* and low *θ* values.

**Fig 9.**
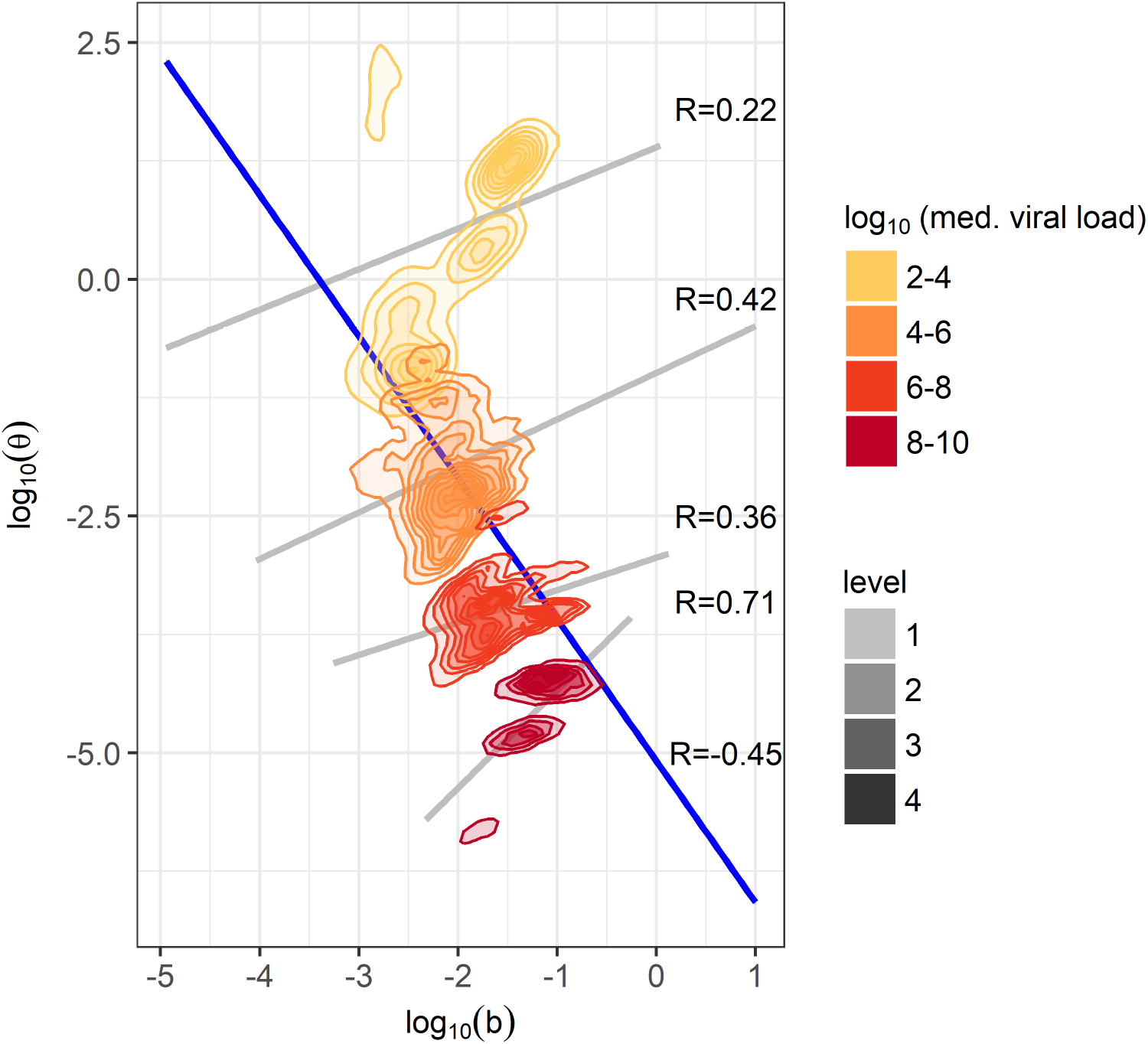
Correlation between parameters *b* and *θ*. Obtained densities of parameters *b* and *θ* are plotted, stratified by median oral EBV viral load group. As *b* and *θ* have opposite effects on viral load, positive correlations are seen within each group (grey lines). However, across all participants, *b* and *θ* are negatively correlated (blue line).

We next examined whether rates of B cell reactivation and cellular immune response could be predicted by the CD4+ T cell count and HIV-1 plasma viral load in HIV-1 co-infected participants. A generalized linear model (Methods) revealed that each log_10_ increase in HIV-1 RNA copies/ml significantly decreased the value of *θ* but did not significantly affect *b*. However, each 100-cell/mm^3^ increase in CD4+ T cell count significantly changed the value of *θ* and *b*, increasing *θ* and decreasing *b* (Table 3).

**Table 3.** Effect of CD4+ T cell count and HIV-1 load on parameters *b* and *θ*. In HIV-1 co-infected participants, median values of parameters *b* and *θ* are influenced by the CD4+ T cell count and HIV-1 plasma viral load. The fold-change (FC) in the fit *b* and *θ* values, for every log_10_ increase in HIV-1 RNA copies/mL and every 100 CD4+ T cell/mm^3^ increase is shown.

|  | Trait | FC | 95% CI | p-value |
| --- | --- | --- | --- | --- |
| $\theta$ | HIV-1 RNA | 0.16 | 0.06-0.42 | <0.001 |
|  | CD4+ T cell | 1.84 | 1.05-3.22 | 0.040 |
| $b$ | HIV-1 RNA | 1.21 | 0.89-1.63 | 0.226 |
|  | CD4+ T cell | 0.80 | 0.69-0.92 | 0.004 |

### Application of our model to a distinct data set from a North American cohort

To test it’s generalizability, we applied our model to a previously described set of data from a cohort of 26 participants in Seattle, Washington, who underwent daily oral EBV sampling and testing using the same methods as the Ugandan cohort described above [22]. A total of 1323 swabs were collected during the 8-week period of the study, with a median of 55 swabs per participant (3-61 swabs). 16 (62%) of these participants were HIV-1 co-infected and if on HAART were required to remain on a stable regimen throughout the study. None were receiving other antiviral drugs at the time of enrollment. While the objective of the Seattle study was to analyse the effects of valganciclovir on daily EBV oral shedding, we restricted our analysis to the data from the eight-week period when participants received placebo.

Participants of the Seattle cohort showed significantly lower oral EBV viral loads than those of the Uganda cohort. Given a positive swab, HIV-1 uninfected, EBV shedding participants had a mean of 1.4 log_10_-lower EBV viral load, while HIV-1 co-infected, EBV shedding participants had a mean of 2.0 log_10_-lower viral load with p-values<0.001 for both cases. Nonetheless, both sets of data serve as good representations of EBV shedding in the saliva and we expect the host-pathogen interactions occurring within the tonsils to be the same. Our mathematical model fit the data again with high fidelity (Fig S3 in Supporting Information) and produced similar results in terms of the number of infected crypts (Fig S4A and S4B in Supporting Information), virus produced by crypts (Fig S4C and S4D in Supporting Information), and the values for parameters *b* and *θ* when stratified by HIV-1 status and median EBV load (Fig S5 in Supporting Information). Full details are given in the Supporting Information. Overall, we find that our modelling approach and findings from the Uganda cohort are strongly supported by analysis of the Seattle cohort data.

## Discussion

By modelling the replication patterns of EBV in saliva in HIV-1 co-infected and uninfected individuals, we were able to evaluate the potential mechanisms that explain why persons with HIV-1 have worse EBV infections and are more susceptible to EBV-related malignancies. Specifically, our model indicates that increased oral EBV shedding with HIV-1 co-infection is due to greater reactivation of EBV-infected B cells as well as impaired EBV-specific cytotoxic T cell immune control.

It has been previously reported that persons infected with HIV-1 have higher oral EBV shedding than HIV-1 uninfected individuals [3, 4, 23–25, 42, 43]. However, few data sets allow detailed representation of the dynamics of oral EBV shedding in HIV-1 co-infected and uninfected participants over time. We also found that HIV-1 infection was associated with a significantly increased frequency and quantity of oral EBV shedding. There was also a statistically significant association between higher CD4+ T cell counts and a lower frequency of EBV shedding in HIV-1 co-infected participants.

B cell activation and plasma cell differentiation have been shown to induce EBV reactivation and lytic replication [12]. Furthermore, it is clear that HIV-1 infection is associated with increased levels of B cell activation [18, 44]. As such, we hypothesized that, in addition to impaired cellular immune control of EBV infection, an increase in EBV reactivation from latently-infected B cells in HIV-1 co-infected individuals contributes to higher levels of oral EBV replication. This prompted us to develop a mathematical model to describe the dynamics of virus, infected cells, and the cellular immune response within the tonsillar epithelium. We then fit this model to participant data, seeking to determine the drivers of these differences.

While previous mathematical models have examined the within-host dynamics of EBV infection [10, 26–29], none have examined the differences between HIV-1 infected and HIV-1 uninfected individuals [38, 45–47]. Strengths of our approach include the incorporation of granular quantitative EBV shedding measurements from two independent cohorts, each made up of HIV-1 co-infected and uninfected individuals, and the availability of CD4+ T cell counts and HIV-1 plasma viral load data from co-infected participants. Limitations include the lack of data on EBV-specific T cell responses and direct measures of B cell activation in these individuals.

Our mathematical model was based on representing the tonsillar epithelium as a series of crypts, each serving as a potential site of epithelial infection and viral shedding. In this way, a single tonsillar crypt behaves similarly to how individual herpes simplex virus (HSV)-2 lesions have been modelled in the past, with reactivation of EBV-infected B cells being analogous to the release of HSV-2 from infected neurons, sparking new epithelial lesions [38, 45–47]. These previous models all captured the stochastic patterns of HSV-2 shedding well, which appear similar to the patterns of oral EBV shedding. Our model assumes that all tonsillar crypts are independent from one another. In reality, virus from one infected crypt may spill over and seed infection in a neighbouring crypt, rather than EBV entering an uninfected crypt purely via the reactivation of B cells as we assume in our model. By not accounting for this, our predicted B cell reactivation rates are likely higher than their true biological values. However, assuming the amount of viral spill-over into neighbouring crypts is proportional to the viral load, our qualitative comparison remains valid. Furthermore, modelling the tonsillar crypts as multiple, segregated sites of infection was essential for simulating the high variability in viral load seen over time in cohort data. In simulations without this spatial separation, viral loads and levels of immune surveillance within an individual often equilibrated over time and no longer matched the stochastic nature of participant data. This result indicates that while virus and immune cells may travel between crypts, this effect is likely minimal. While a fully-spatial model including the mobility of virus and immune cells throughout the tonsils would be ideal, it could only be parameterized speculatively, and would be computationally intensive. Our intermediate strategy of a crypt-level simulation is simple enough to be computationally feasible but complex enough to retain the inherent stochasticity and spatial diversity of EBV infection dynamics within the tonsils.

Using our mathematical model, we found that HIV-1 co-infected individuals are expected to have more actively infected crypts producing more EBV at any given time than HIV-1 uninfected individuals. Also, by specifically including the dynamics of cellular immune control and rates of B cell reactivation leading to new epithelial infection, we saw that both factors contribute to higher EBV viral loads in HIV-1 co-infected individuals. These results are consistent with previous observations that cellular immune control of EBV infection in HIV-1 co-infected individuals is impaired [48]. Similarly, HIV-1 infection is known to dysregulate B cell function and increase B cell activation, which can result directly from HIV-1 itself as well as more frequent and severe co-infections compared to HIV-1-uninfected individuals [17, 18, 49, 50]. Indeed, our model demonstrated that higher CD4+ T cell counts were associated with significantly higher predicted rates of EBV-specific cytotoxic T cell activity and significantly lower predicted rates of EBV-infected B cell reactivation. Furthermore, increases in HIV-1 plasma RNA levels were associated with significantly lower predicted rates of EBV-specific cytotoxic T cell activity.

Importantly, our results have implications for strategies to prevent EBV infection and disease. EBV-specific cellular immunity is recognized as critical for controlling EBV replication and preventing EBV-associated malignancies. [48, 51–55]. Independent of restoring EBV-specific cellular immune responses, strategies to reduce B cell reactivation in EBV-infected persons might limit viral replication, transmission, and related malignancies [56, 57].

## Methods

### Cohorts and samples

Men and women aged 18 to 65 were enrolled in the Uganda cohort as previously described and were followed for four weeks [31]. Eligible HIV-1 seropositive persons had a CD4+ T cell count greater than 200 cells/mm^3^ and were not taking antiretroviral therapy, in accordance with the WHO guidelines at that time [58]. Men aged 24 to 66 were enrolled in the Seattle cohort as previously described [21, 22].As the Seattle cohort shedding data was obtianed from a randomized placebo-controlled cross-over trial of valgnaciclovir, only data collected while participants were receiving placebo was used for this study. Both participants and pill administrators were unaware of group assignments. Participants of both cohorts did not take any drugs with anti-herpesvirus activity during the study period. Self-collected daily oropharyngeal swabs for both cohorts were collected by swabbing the oral mucosa and pharynx with a Dacron swab and were then placed in a vial containing 1 ml of 1X digestion buffer, stored at room temperature, and returned at weekly (Ugandan cohort) or bi-weekly (Seattle cohort) clinical visits. In the Uganda cohort, focused physical exams and collection of genital and plasma samples were performed at weekly clinic visits. This data is described in the Supporting Information. All participants provided either written or verbal informed consent. Study procedures for both cohorts were approved by the University of Washington Human Subjects Review Board. Additional approval for the Ugandan cohort study was given by the Makerere University Research and Ethics Committee and the Uganda National Council for Science and Technology.

### Laboratory testing

Commercially available immunoassays were used to ascertain HIV-1 and EBV serostatus (Inverness Medical Innovations, Inc and Wampole^®^ for the Ugandan cohort and Abbott Laboratories for the Seattle Cohort [22]). For the Ugandan cohort, CD4+ T cell counts, and plasma HIV-1 RNA levels were determined in HIV-1-infected persons at the Makerere University-John Hopkins University laboratory using standard cell sorting techniques and the Amplicor HIV-1 monitor test (Roche, version 1.5), respectively. For both cohorts, DNA was extracted from mucosal swabs and plasma [59], and real-time quantitative polymerase chain reaction (qPCR) was performed using specific primers to detect EBV [60], with positive and negative controls as previously described [19, 59]. Mucosal samples with greater than 150 copies/ml and plasma samples with greater than 50 copies/ml herpesvirus DNA/ml were considered positive [61].

### Statistical analyses of data

The frequency of mucosal shedding and viremia was defined as the proportion of samples testing positive for EBV. The frequency of mucosal shedding was first compared in HIV-1 co-infected and uninfected persons. To do this, we used generalized estimating equations (GEE) and assumed frequencies followed a Poisson distribution. In HIV-1 seropositive persons, frequencies of mucosal shedding and viremia were also modelled with GEE allowing for continuous adjustment for each 100 cell/mm^3^ increase in CD4+ T cell count and each log_10_ increase in HIV-1 RNA. Again, we assumed the frequency of shedding follows a Poisson distribution. Finally, GEE were used to compare the quantities of virus shed in mucosal samples in HIV-1 co-infected and HIV-1 uninfected persons assuming a Gaussian distribution. In all tests, two-sided p-values ≤0.05 were considered statistically significant.

### Mathematical model simulations

Based on the reactions of the model, we applied the tau leaping algorithm to stochastically simulate the dynamics of each crypt [62]. With this algorithm, a small, constant-sized time step is taken, and the number of occurrences of each reaction is stochastically chosen following a Poisson or Multinomial distribution depending on the independence of the reaction. One long simulation is performed, which is then divided into 240 sections to represent the dynamics of each of the 240 crypts. Specifically, the simulation is run out to a time of

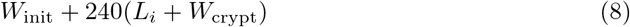

and crypt *c*’s dynamics are taken from the time interval

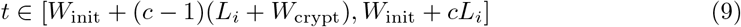

where *W*_init_ represents the time necessary to remove the effects of the initial conditions on the simulation, *L_i_* represents the duration of which participant *i* had oral swabs taken, and *W*_crypt_ represents the time necessary to make the dynamics of one crypt quasi-independent of the next. With immune cell decay (*δT*) acting as the slowest rate in the model (*δ* = 0.1/(day-cell)), we let both *W*_crypt_ and *W*_init_ equal 120 days, so that if infection in one crypt occurred, only an expected 1/10^6^ of the responding immune cells would carry on to the next crypt’s dynamics.

Viral loads from each crypt are added together to get the model-predicted amount of virus seen in the saliva over time. As the qPCR threshold of detection was 150 copies/ml for the data used, whenever the total simulated viral load in the saliva dropped below 150 copies/ml, we set the output to zero.

### Model fitting using Approximate Bayesian Computation

We next fit parameters to daily quantitative oral EBV shedding data from our cohort participants. We used Approximate Bayesian Computation (ABC), where summary statistics of the data and model simulations are compared to determine which parameters allow the model to best fit the data. We used the R package EasyABC to execute sequential ABC, following Lenormand’s algorithm [63]. Here, uniform priors for each parameter are set, and an initial *n* number of simulations are run, each with a different set of parameters randomly chosen from the priors. The algorithm calculates summary statistics, chosen by the user, for each parameter set 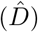 and compares how well they match with the summary statistics of the data (*D*) by calculating a distance measure, 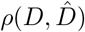. The best-matching *ϕ* percent of these simulations are kept, with the parameters of the chosen simulations used to build new priors. This process repeats until the distance between the summary statistics of the data and simulations is minimized. We executed this algorithm for the data of each individual participant. We chose to capture the trends of the data using 5 summary statistics: the frequency of positive swabs, the median, maximum, and variance of detectable viral loads, and the number of peaks in viral loads, a peak being defined as when the directly preceding and following time points have lower viral loads. The associated *ρ* value for participant *i* and parameter set *j* (*ρ_i,j_*) is defined as in Equation 7 of the main text:

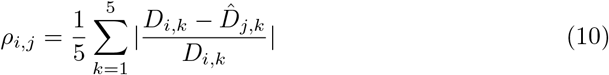

where *D_i,k_* is the *k^th^* summary statistic for the data of participant *i* and 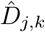 is the *k^th^* summary statistic for parameter set *j*. Using the Lenormand algorithm, 1000 parameter sets that minimize *ρ_i,j_* were selected for each participant. During this fitting, we fixed all parameters except *b* and *θ* as these are the two parameters that are likely most affected by HIV-1 infection. For each chosen parameter set, we also calculated the median number of actively infected crypts and the median amount of virus produced by an actively infected crypt, so we could later compare how these traits differ between participant groups.

Since we lack information on immune cell presence in tonsillar crypts and can only fit the model to data on viral load, we had to censor simulations where the cytotoxic T cell level became unrealistically high. Whenever a parameter set led to a simulation where cytotoxic T cell count within a tonsillar crypt was greater than 10^5^ cells, we prevented it from being selected by the ABC algorithm, ensuring only biologically relevant simulations were being considered.

### Determining the posterior distribution of parameters and differences between participant groups

We combined the results of our ABC fitting algorithm to compare how the posterior distributions of parameters *b* and *θ*, the number of actively infected crypts, and the amount of virus produced vary between different participant groups.

As some parameter values selected by the ABC fitting algorithm fit the data better than others (i.e. have lower *ρ* values), we approximated the posterior distributions of our parameter sets by performing importance sampling on the raw posterior distributions [64, 65]. To do this, we weighted each output parameter set by the reciprocal of its *ρ* value. By weighting inversely to *ρ*, we assume our model is a correct representation of viral dynamics in the tonsils and put greater importance on those parameter sets that fit the data well.

To determine the posterior distributions of parameters in HIV-1 co-infected and uninfected groups (*X_A_* and *X_B_* respectively), the probability of each parameter set (*x*(*i,j*)) serving in each posterior is set to

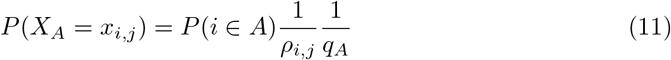

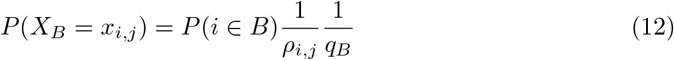

where *A* and *B* are the sets of indices of participants who are HIV-1 uninfected and co-infected, respectively, and we define the normalization factors

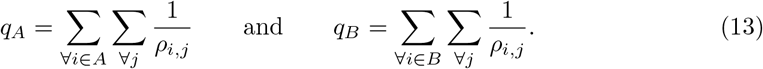

Note that

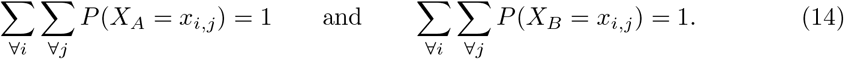

We took 10^5^ draws from each distribution and plotted the resulting data to obtain graphical representations of the posterior parameter distributions for parameters *b* and *θ*, the number of actively infected crypts, and the virus produced, for HIV-1 co-infected and uninfected participants. The above process was repeated where instead of stratifying by HIV-1 status, participants were stratified by median EBV load in order to produce similar plots.

We also performed importance sampling on the raw posterior distributions for individual participants. Using these, we were able to calculate the mean parameter values for *b* and *θ* for each participant. Means of parameters *b* and *θ* in HIV-1 co-infected participants were then modelled as functions of the participants’ CD4+ T cell count and HIV-1 RNA load. This was done using GLM allowing for continuous adjustment for each 100 cell/mm^3^ increase in CD4+ T cell count and each log_10_ increase in HIV-1 RNA. Parameters *b* and *θ* were assumed to follow a Gaussian distribution. In these tests, two-sided p-values ≤0.05 were considered statistically significant.

### Sensitivity analysis of model parameters

We performed two sensitivity analyses to evaluate how changes in parameter values affect the results of the model. First, to initially determine acceptable values for parameters, we performed a univariate analysis, starting with an initial set of parameters, varying one parameter at a time, and running 100 simulations for each parameter set to gain a representation of how EBV viral dynamics behave. From this analysis we selected parameter values for *β, f, α, δ, p*, and *c* which would remain fixed throughout the ABC data fitting process while leaving the parameters of most interest, *b* and *θ*, free.

After completing ABC, we performed another univariate analysis to observe whether our choices for fixed parameter values were correct. By letting *b* and *θ* equal values selected by the ABC algorithm and then individually varying the parameters that were fixed, we checked whether different values of our fixed parameters would have improved the model’s fit.

