## Supplementary information for "Increased oral Epstein-Barr virus shedding with HIV-1 co-infection is due to a combination of B cell activation and impaired cellular immune control"

### Supporting information

#### Ugandan Cohort demographics

Within the Ugandan cohort, all 85 participants provided at least one oral swab and were included in this analysis. 32 participants (38%) were female and 53 (62%) were male with a median age of 32 years (range 18-60 years). Of the 42 participants (49%) who were HIV-1 seronegative, 20 (48%) had KS. All individuals who tested positive for HIV-1 also provided blood to determine CD4+ T cell count and HIV-1 RNA levels at a single time point. The median CD4+ T cell count among participants was 439 cells/mm$^3$ (range 208-1062 cells/mm$^3$) while the median HIV-1 RNA load among participants was 4.7 $\log_{10}$ copies/ml (range 2.3-5.9 $\log_{10}$ copes/ml). Serum was also provided by 82 of the 85 participants to determine EBV serostatus; all tested positive. The remaining 3 participants had confirmed EBV infection based on the direct detection of virus in oral swabs. Therefore, all participants in this study had evidence of EBV infection. Additional details of this cohort have been published in [1].

#### Detection of EBV within Ugandan participants' oral and genital swabs, and plasma samples

Ugandan cohort participants had daily oral swabs, weekly genital swabs and plasma samples taken to test for EBV. The percentage of oral swabs testing positive for EBV stratified by HIV-1 status is displayed in Fig 1A and was used in our mathematical model, while the percentage of genital swabs and plasma samples testing positive for EBV are shown in Fig S1A. All (100%) HIV-1 seropositive participants had at least 1 EBV positive oral swab, while 40 out of 42 (95%) HIV-1 seronegative participants had at least 1 EBV positive oral swab. 1030 of the 1093 (94%) oral swabs taken from HIV-1 seropositive participants tested positive for EBV while 867 of the 1171 (74%) oral swabs taken from HIV-1 seronegative participants tested positive for EBV. Of the participants who provided genital swabs, 31 of the 41 (76%) contributing HIV-1 seropositive participants had at least 1 EBV positive genital swab, while 13 of the 38 (34%) contributing HIV-1 seronegative participants had at least 1 EBV positive genital swab. 71 of the 154 (46%) genital swabs taken from HIV-1 seropositive participants tested positive for EBV while 30 of the 139 (22%) genital swabs taken from HIV-1 seronegative participants tested positive for EBV. EBV was detected in plasma from 30 of 42 (71%) HIV-1 seropositive participants and 19 of 42 (45%) HIV-1 seronegative participants. 60 of the 208 (29%) plasma samples from HIV-1 seropositive participants tested positive for EBV compared to 41 of the 218 (19%) plasma samples from HIV-1 seronegative participants.

#### Association between participant HIV infection and CD4+ T cell characteristics and the EBV loads in genital swabs and plasma samples

While the determinants of EBV loads in the saliva are analysed in the main paper, the effects of HIV-1 status, HIV-1 load and CD4+ T cell count on the EBV loads in Ugandan participants' genital swabs and plasma samples are analysed here. Descriptions of the fraction of participant swabs testing positive for EBV, and the viral loads within positive swabs, are shown in Fig S1. These data were next used to determine whether there was a significant relationship between the frequency of viral shedding at these sites and the HIV-1 status of participants, or the CD4+ T cell count or HIV-1 load in HIV-1 co-infected participants. Results are shown in Table S1 and Table S2. Participants who were HIV-1 infected had significantly higher frequencies of EBV detection in genital swabs than participants who were HIV-1 uninfected (p-value of 0.007); however, this was not true for plasma samples (p-value of 0.103). When examining the median amount of virus shed by each participant, HIV-1 infection status showed no significant effect (p-values of 0.626 and 0.080 for genital and plasma samples respectively, Fig S1B). In those who were HIV-1 co-infected, CD4+ T cell count significantly decreased the frequency of EBV detection in the plasma but not in genital swabs (p-values of 0.029 and 0.137 respectively). Increases in the amount of HIV-1 RNA detected had no significant effect on the frequency of EBV detection at either site.

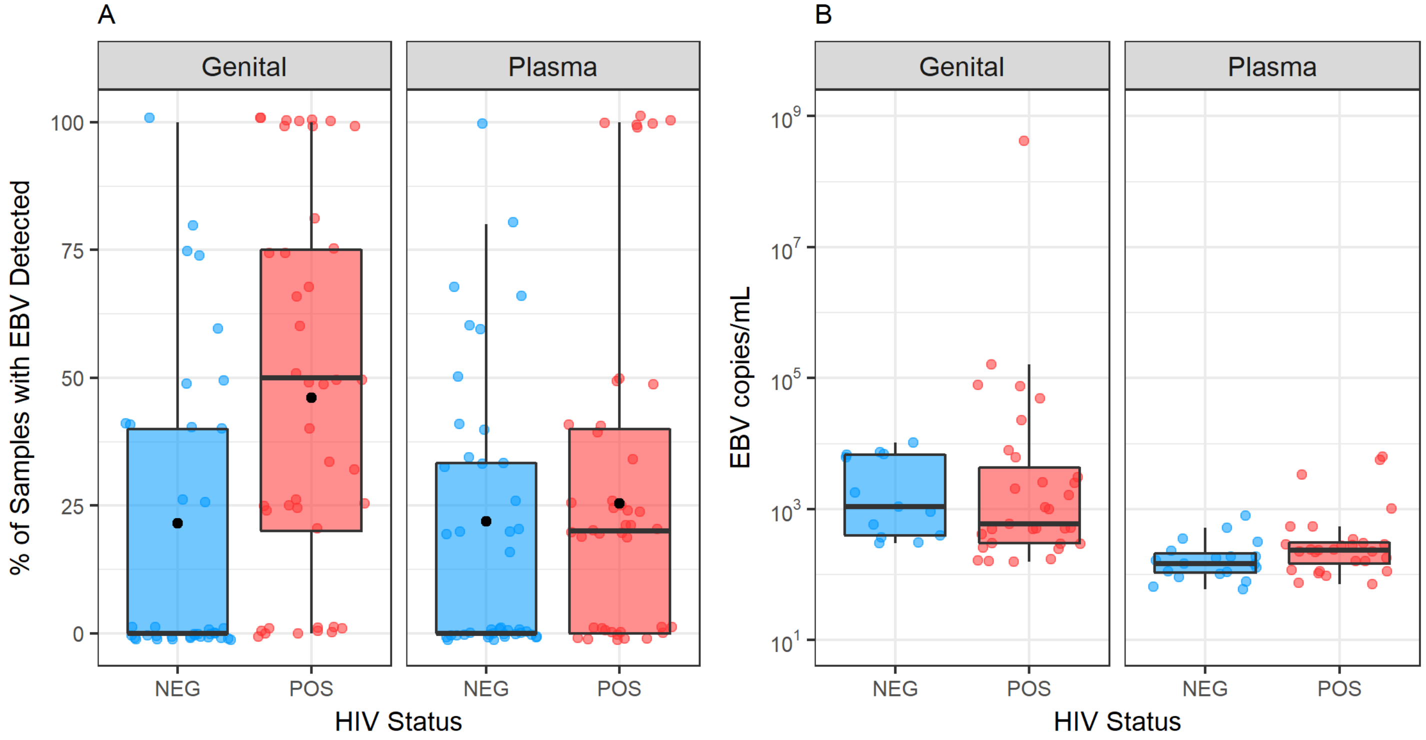


**Fig S1. Impact of HIV-1 infection on EBV detection in Ugandan genital swabs and plasma samples.** A. Percentages of samples that tested positive for EBV for each participant stratified by HIV-1 status and collection site. Black dots indicate the percentage of samples that tested positive for EBV when pooling participant samples. B. Median EBV viral loads/ml in oral swabs testing positive for EBV, per participant stratified by HIV-1 status and collection site.

**Table S1. Effect of HIV-1 seropositivity on the frequency of EBV detection in genital swabs and plasma samples.** The incidence rate ratio (IRR), 95% confidence interval (CI) and p-value are shown for genital swabs and plasma samples.

| | **IRR** | **95% CI** | **p-value** |
|---|---|---|---|
| **Genital** | 2.14 | 1.23-3.71 | 0.007 |
| **Plasma** | 1.53 | 0.92-2.56 | 0.103 |

**Table S2. Effects of plasma HIV-1 load and CD4+ T cell count on the frequency of EBV detection in genital swabs and plasma samples.** The effects of each 100-cell increase in CD4+ T cell count and each $\log_{10}$ increase in HIV-1 RNA in the serum on the frequency of EBV detection and their p-values are shown.

| | **Trait** | **IRR** | **95% CI** | **p-value** |
|---|---|---|---|---|
| **Genital** | HIV-1 RNA | 0.92 | 0.77-1.11 | 0.393 |
| | CD4+ T cell | 0.90 | 0.79-1.03 | 0.137 |
| **Plasma** | HIV-1 RNA | 1.55 | 0.95-2.53 | 0.079 |
| | CD4+ T cell | 0.85 | 0.73-0.98 | 0.029 |

#### Validation of the mathematical model using a North American cohort

To observe how well our model performs when applied to other cohorts, we used data from another previously described cohort where daily EBV shedding in the oral mucosa was measured in 26 adult men from Seattle, Washington [2]. The distribution of viral loads found in positive swabs for each participant is shown in Fig S2. Viral loads in this cohort were lower than those seen in our Uganda cohort: HIV-1 uninfected participants had a mean of 1.2 $\log_{10}$ lower viral loads, and HIV-1 co-infected participants had a mean of 1.9 $\log_{10}$ lower viral loads, with p-values<0.001 for both. However, both data sets serve as good representations of EBV shedding in the tonsils and all individuals should have similar viral dynamics that can be described by our mathematical model.

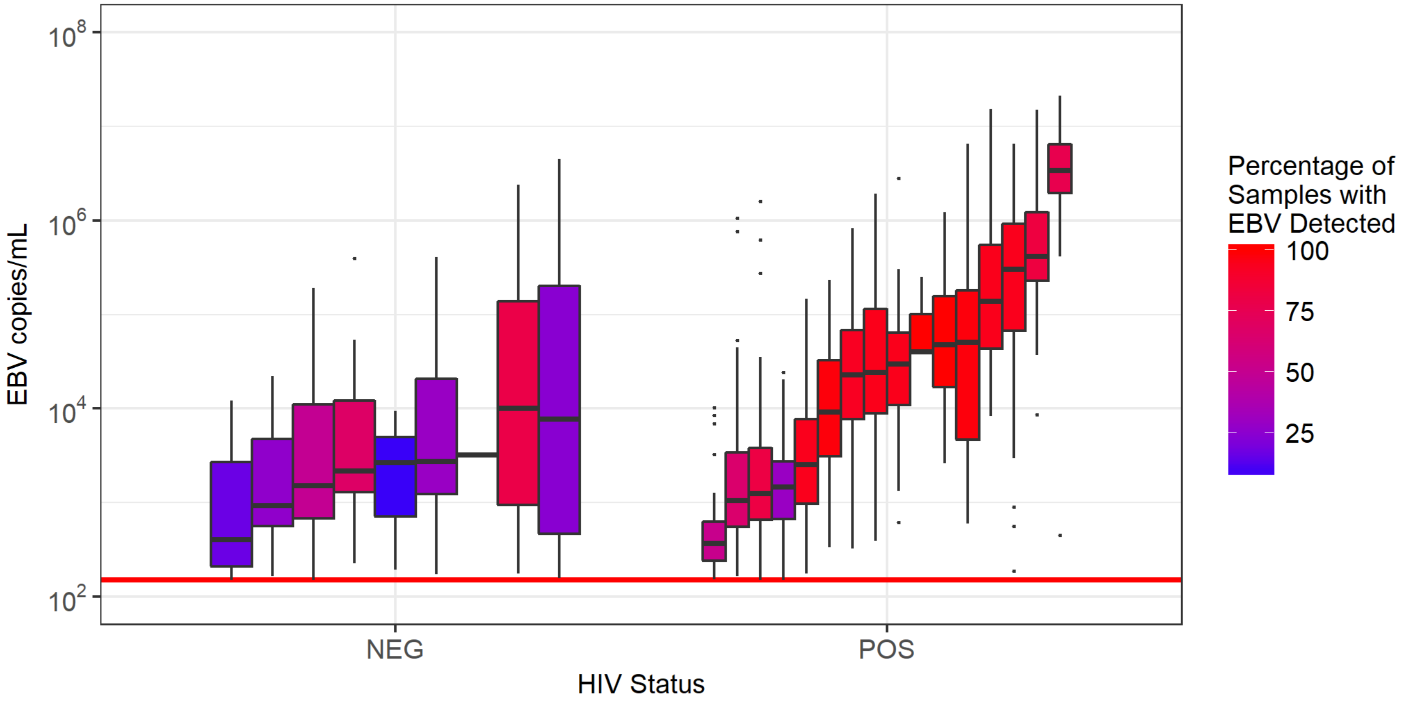


**Fig S2. Distribution of virus loads detected in oral swabs from participants of the Seattle cohort.** Each box and whisker represents the distribution of the viral loads in daily oral swabs testing positive for EBV for an individual participant. Since many swabs did not test positive for EBV via qPCR, the percentage of a participant's oral samples that did test positive is indicated by the colour of the box. The threshold of detection (150 copies/ml) is indicated by the horizontal red line.

We applied the ABC fitting algorithm as described in the Methods to fit individual participant data from the Seattle Study to our model. We were able to determine parameter fits for 23 of the 26 participants. Those individuals whose data could not be fit either had no EBV viral shedding detected (1 participant), only 1 positive swab for EBV (1 participant) or an insufficient number of data points (only three swabs for 1 participant). The distribution of $\rho$ for each participant is shown in Fig S3. We found that the $\rho$ values are similar for participants in both studies. While the mean $\rho$ value was significantly lower ($p<0.001$) for participants in our Uganda study (mean of 0.19) than for participants in the Seattle study (mean of 0.33), this was mainly due to the model fitting better to individuals with higher viral loads (Results). When continuously correcting for unit increases in the $\log_{10}$ of the participants' median EBV load, there was no significant difference between $\rho$ values from Ugandan participants and the participants in the Seattle cohort (mean increase in $\rho$ of 0.05 for Seattle study compared to our study, 95% CI = [-0.01,0.11], p-value = 0.096).

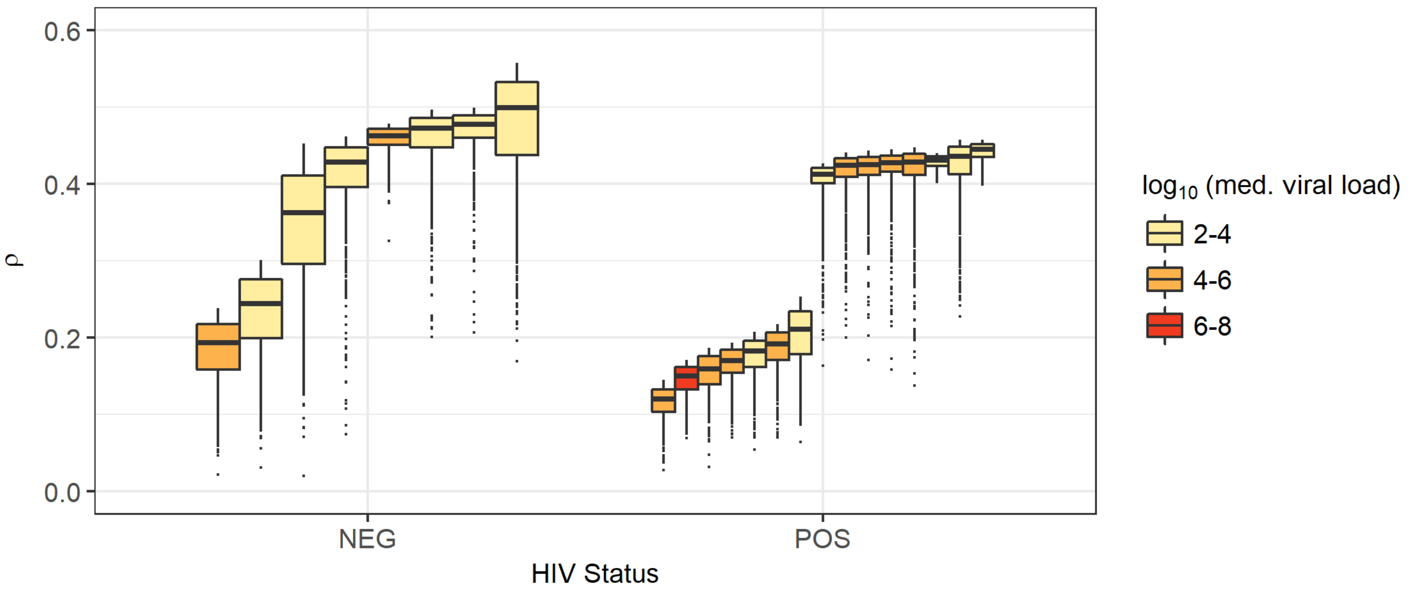


**Fig S3. Distribution of goodness of fit ($\rho$) statistics for participants of the Seattle study.** The data from 23 of the 26 Seattle study participants were able to fit to our model. For each participant, the 1000 parameter sets that produced the best agreeance between the summary statistics of the data and model simulations were chosen. The $\rho$ values, a measure of the fit, is shown for each parameter set. $\rho$ values for participants of the Seattle study are similar to those for participants of our study, indicating the model works equally well for both sets of data.

As described in the Methods, we then used importance sampling to determine distributions for the parameters governing infection traits in different participant groups. Specifically, we looked at the median number of tonsillar crypts actively infected in an individual (Fig S4A and S4B), the median amount of virus within an active crypt (Fig S4C and S4D), the rate or B cell reactivation leading to epithelial infection ($b$) (Fig S5A and S5B), and the rate at which EBV-specific cytotoxic T cells, effective at clearing infection, proliferate in response to infection (Fig S5C and S5D).

Since the Seattle study participants had significantly lower viral loads than participants of our study, it is unsurprising that they had a lower median number of crypts actively producing virus at any given time. Our model predicts that HIV-1 uninfected and HIV-1 co-infected individuals both have a median of 0 crypts actively infected, with a probability of only 0.37 that the number of crypts actively infected in an HIV-1 co-infected individual is greater than the number of crypts actively infected in an HIV-1 uninfected individual. We previously observed from the Uganda cohort that the median viral load of the individual is positively correlated with the number of actively infected crypts at any given time. This remains true for the Seattle study (Fig S4B) although the correlation is not as strong. Similar results appear when looking at the median amount of virus within an actively infected crypt. At any given time, we estimate that the virus load within an actively infected crypt is greater for HIV-1 co-infected individuals with probability 0.58 (compared to HIV-1 uninfected individuals). We also find a positive correlation between the EBV load of an individual and the expected amount of virus that is produced by each of their actively infected crypts. When forming predictions based on the Seattle study data, an individual with 4-6 $\log_{10}$ EBV DNA copies/ml has a higher median amount of virus within a single actively infected crypt than that of an individual with 2-4 $\log_{10}$ EBV DNA copies/ml with probability 0.88, and individual with 6-8 $\log_{10}$ EBV DNA copies/ml has a higher median amount of virus within a single actively infected crypt that that of an individual with 4-6 $\log_{10}$ EBV DNA copies/ml with probability 0.76.

When evaluating differences between the densities of parameters $b$ and $\theta$ in individuals of different HIV-1 statuses and with different median EBV loads, the Seattle study again gives similar predictions to those achieved with our data. From this data we can predict that parameter $b$ is higher in an HIV-1 co-infected individual than an HIV-1 uninfected individual with probability 0.87, and that parameter $\theta$ is lower in an HIV-1 co-infected individual than in an HIV-1 uninfected individual with probability 0.65. These probabilities were similar (0.76 and 0.74 respectively) in the Uganda cohort. When comparing $b$ and $\theta$ values across individuals with different median EBV loads we see that Seattle study individuals with a median EBV load of 4-6 $\log_{10}$ copies/ml in their saliva have a higher value for parameter $b$ and/or $\theta$ than individuals with a median EBV load of 2-4 $\log_{10}$ copies/ml with probability 0.65 and 0.93 respectively, and individuals with a median EBV load of 6-8 $\log_{10}$ copies/ml in their saliva have a higher value for parameter $b$ and/or $\theta$ than individuals with a median EBV load of 4-6 $\log_{10}$ copies/ml with probability 0.80 and 0.98 respectively.

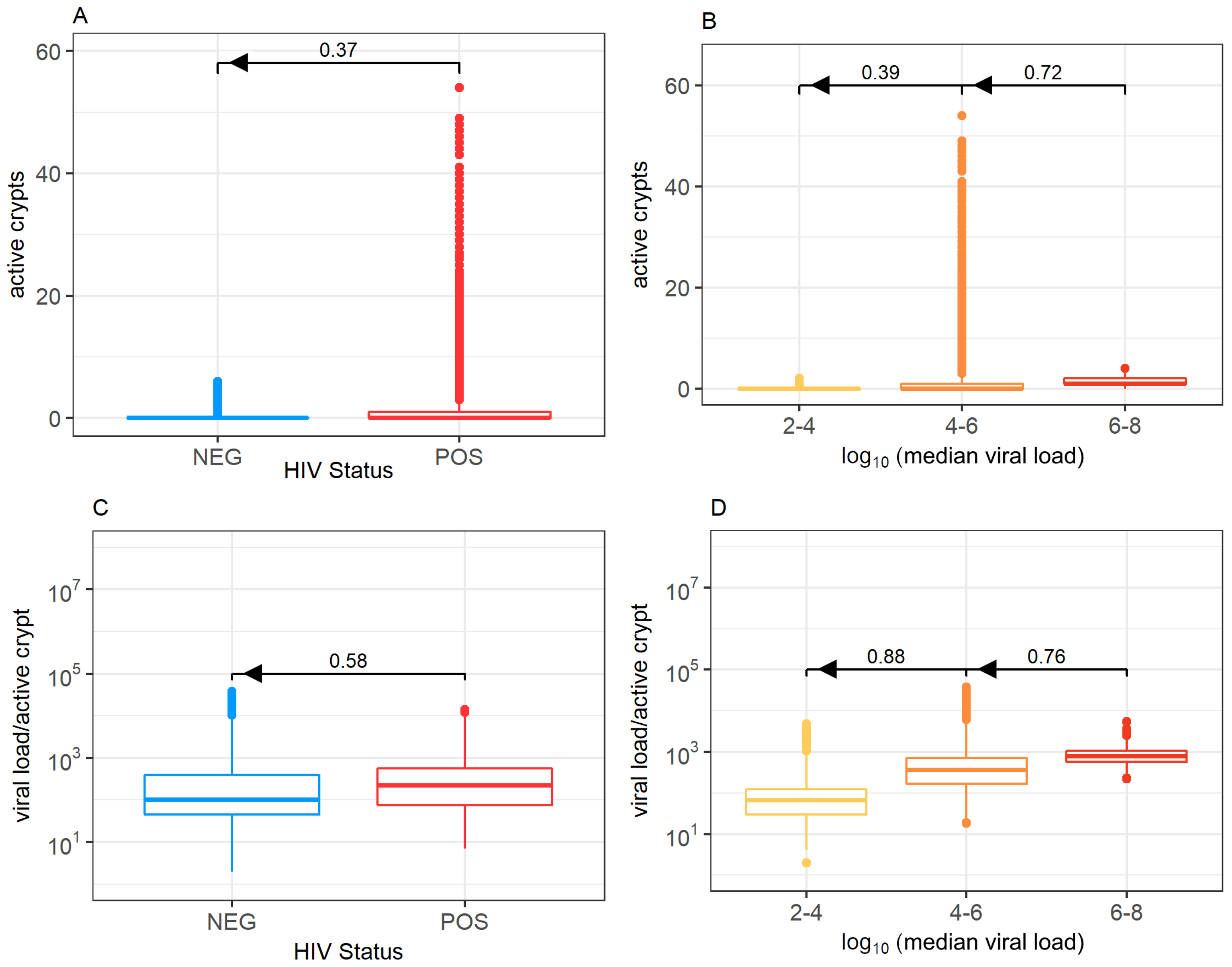


**Fig S4. Numbers of actively infected crypts and EBV viral loads from the Seattle study.** We present distributions of the median number of crypts actively producing EBV within an individual, stratified by (A) HIV-1 status and (B) the $\log_{10}$ median viral load of the individual. We also show distributions of the median amount of virus within a crypt actively producing virus stratified by (C) HIV-1 status and (D) $\log_{10}$ median viral load. Bars above boxplots indicate the probability that a randomly selected individual of one group has a higher parameter value (number of active crypts or viral load per active crypt) than a randomly selected individual in a second group. Arrows show the direction of comparison.

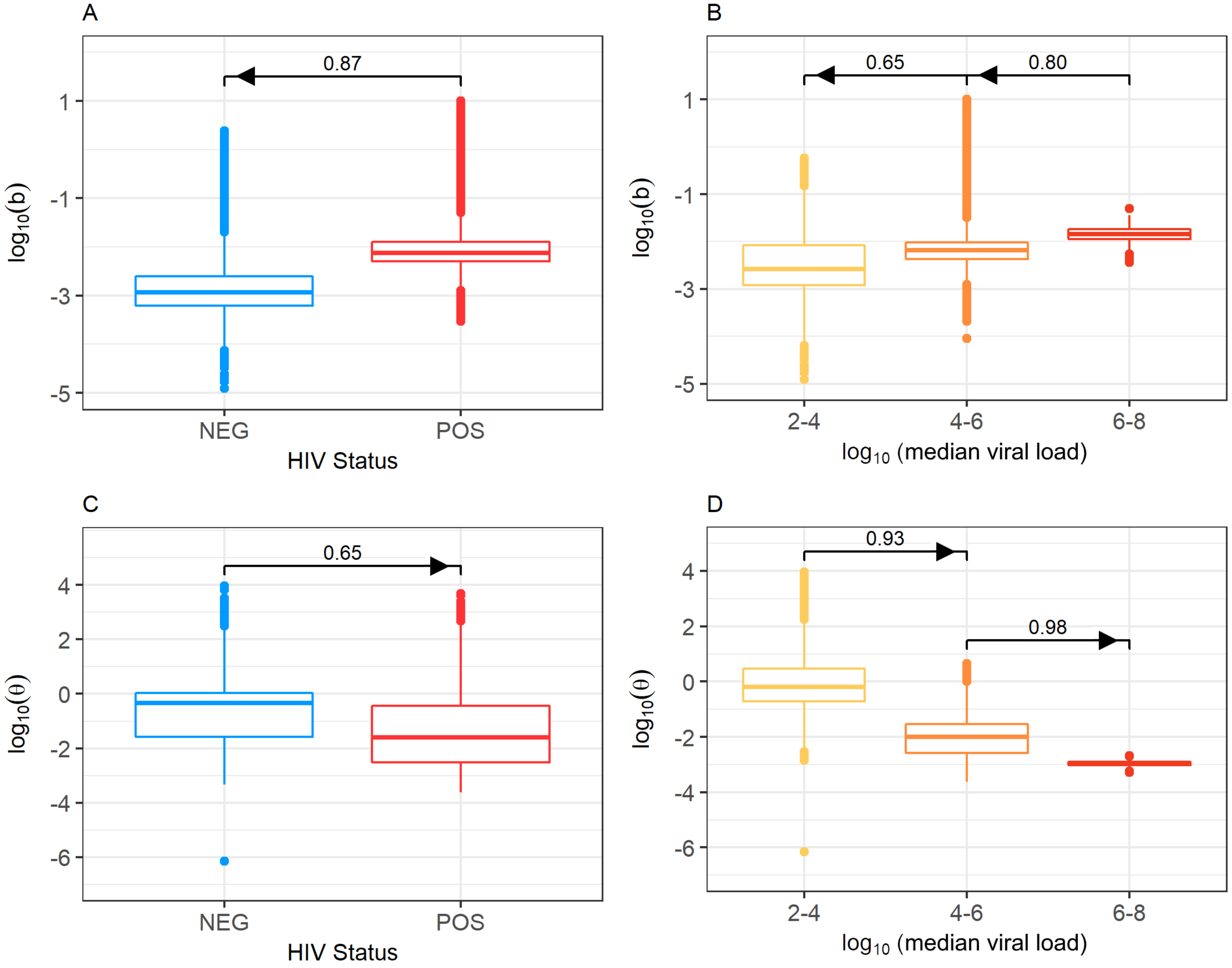


**Fig S5. Distribution of parameters $b$ and $\theta$, stratified by HIV-1 status and median EBV viral load in the Seattle study.** Fitting our mathematical model to participant data revealed that parameter $b$ is greater in HIV-1 co-infected participants (A), increasing with median viral load (B), and parameter $\theta$ is lower in HIV-1 co-infected participants (C), decreasing with median viral load (D). Bars above boxplots indicate the probability that a randomly selected individual of one group has a higher parameter value (be it $b$ or $\theta$) than a randomly selected individual in a second group. Arrows show the direction of comparison.

The Seattle study also collected data on CD4+ T cell count and HIV-1 RNA load in the plasma of HIV-1 co-infected participants. We sought to understand how this data correlated with our predicted values of parameters $b$ and $\theta$. We performed importance sampling on the parameter sets chosen for each participant based on the corresponding goodness of fit ($\rho$) values to find the posterior distribution of $b$ and $\theta$ for each participant. We then took the median value for each participant, and using GLM, determined how each 100-cell increase in CD4+ T cell count and each $\log_{10}$ increase in HIV-1 RNA load affected these values. Results are shown in Table S3. While these traits did not cause significant changes to $b$ or $\theta$, they did cause changes in the same direction as those found when performing the same analysis on the Uganda cohort data.

The lack of significance may be due to the smaller cohort size of the Seattle study.

**Table S3. Effect of CD4+ T cell count and HIV-1 load on the median value of parameter $b$ and $\theta$ for individuals in the Seattle study.** In HIV-1 co-infected participants, median values of parameters $b$ and $\theta$ are influenced by the CD4+ T cell count and HIV-1 load. Fold-change is shown.

| | **Trait** | **FC** | **95% CI** | **p-value** |
|---|---|---|---|---|
| $\theta$ | HIV-1 RNA | 0.33 | 0.12-0.95 | 0.060 |
| | CD4+ T cell | 0.96 | 0.52-1.75 | 0.889 |
| $b$ | HIV-1 RNA | 1.00 | 0.66-1.50 | 0.991 |
| | CD4+ T cell | 0.91 | 0.74-1.11 | 0.351 |

Overall, we find that our mathematical modelling approach performed well in fitting the data from the Kampala, Uganda cohort and from the Seattle, Washington study, validating its use as a reliable description of EBV infection dynamics in the tonsils. We also found quantitatively similar results concerning the effects of HIV-1 co-infection among both sets of participants.
